# From APTs to advanced persistent biological threats (APBTs): a taxonomy and large-scale metadata audit of genomic surveillance infrastructure

**DOI:** 10.64898/2026.10.09.757979

**Authors:** Nasreen Anjum, Mahreen Kiran, Hessa Abdulrahman Alfraihi, Aikaterini Kanta, Mahreen ul-Hassan, Brandon Jeffery Weihs

## Abstract

Open genomic repositories have become critical infrastructure for pathogen surveillance, variant tracking, outbreak reconstruction, and public health decision making. However, many of these repositories were designed to support rapid scientific data sharing rather than adversarial verification, creating potential security gaps. This paper introduces Advanced Persistent Biological Threats (APBTs), as a framework for understanding how malicious cyber biological actors can gradually disrupt genomic surveillance without entering a laboratory or releasing a biological agent. Instead of altering biological material, an attacker targets the sequence data, metadata, reference datasets, and analytical models used by surveillance systems. We present, to our knowledge, the first integrated multilayer taxonomy of APBTs, structured around a four phase lifecycle, and investigate whether the metadata weaknesses identified by this framework are present in an operational genomic repository. To assess this, we audited 100,000 SARS-CoV-2 BioSample records obtained from the NCBI. A framework of 23 checks was applied across five metadata layers: temporal, geographic, host and specimen, provenance, and technical metadata. Every record triggered at least one APBT relevant metadata vulnerability indicator, with a mean of 7.34 indicators per record (SD = 1.34). Overall, 97.3% of records were classified in the High or Critical severity categories. Every record also contained at least one indicator in both the provenance and technical metadata layers. These results largely reflect the optional status of several fields in the current BioSample submission model rather than isolated errors by individual data contributors. Temporal and geographic information was comparatively complete, while fields needed for provenance reconstruction, chain of custody assessment, technical validation, and analytical interpretation were frequently missing. These findings do not indicate deliberate manipulation. However, the structural incompleteness of BioSample metadata creates vulnerabilities that an adversary can exploit to introduce poisoned, misleading, or weakly traceable metadata into genomic repositories. As a result, maliciously altered metadata can appear plausible, remain undetected, and influence downstream genomic analysis and surveillance. The findings therefore show that widely used pathogen genomic repositories already contain structural weaknesses that can support metadata poisoning as part of an APBT operation.

## 1 Introduction

Genomic data science has become an important component of modern public health. Its importance became especially evident during the COVID-19 pandemic. Within weeks of the first reported cases, the SARS-CoV-2 genome had been sequenced and shared internationally, supporting the rapid development of diagnostic tests, vaccines, and variant surveillance systems [1–3]. As sequencing capacity expanded, global genomic repositories and data-sharing platforms, including NCBI GenBank, the European Nucleotide Archive (ENA), and GISAID, accumulated millions of pathogen genomes. These resources enabled researchers and public health agencies to track variant emergence, examine transmission dynamics, and support outbreak response and public health decision making in near real time [4–6]. These developments positioned genomic surveillance as both a means of detecting biological change and a critical component of global public health information infrastructure.

The openness and automation that give genomic surveillance its speed and scale also expose these systems to data integrity risks. The reliability of genomic sequences, associated metadata, reference datasets, and machine learning (ML) outputs depends on the integrity, consistency, and traceability of the underlying data. However, the repositories and analytical pipelines that manage these resources were developed primarily to support scientific collaboration, accessibility, and rapid data sharing, with less emphasis on resilience against deliberate manipulation. Previous research has documented several distinct categories of integrity risk, including inconsistencies in repository metadata [7], weaknesses in the screening of designed or synthetic sequences [8], and the sensitivity of biomedical ML systems to biased or corrupted training data [9]. These risks have largely been examined as separate technical problems rather than as interconnected components of a persistent adversarial campaign affecting genomic surveillance reliability.

This paper introduces Advanced Persistent Biological Threats (APBTs) as a threat concept for biological surveillance systems, analogous to Advanced Persistent Threats (APTs) in cybersecurity. Conventional APTs target computational infrastructure, including networks, servers, software, and data systems [10, 11]. In contrast, APBTs target the informational infrastructure of public health, including genomic sequences, metadata, reference datasets, analytical models, and governance processes used in biosurveillance. Like APTs, APBTs involve long-term planning, gradual and individually inconspicuous actions, and strategic objectives that remain concealed until an opportune moment. However, they do not necessarily require direct system intrusion or the manipulation of biological materials. Instead, an operation can be conducted through the routine data submission processes of open scientific repositories.

By targeting the information used in genomic surveillance, APBTs can reduce the reliability of subsequent analyses and decisions. Their strategic significance has two main dimensions. First, the gradual corruption of genomic data can delay variant detection, distort the understanding of pathogen evolution and transmission, reduce the accuracy of epidemiological models, and contribute to inconsistent public health responses [12, 13]. Second, APBTs can weaken surveillance capacity before a major biological event. If a new pathogen later emerges or is deliberately released, a compromised system may be slower to detect the event, assess its characteristics, and determine its likely source.

Previous research has examined genomic repository integrity, metadata provenance, AI-enabled sequence generation, data poisoning, and the security of genomic analysis systems. However, these topics have largely been studied in isolation. Bringing them together is necessary to understand how coordinated and persistent manipulation can affect genomic surveillance across the data lifecycle. The following section reviews these areas, identifies the remaining research gap, and positions this study as an empirical assessment of APBT-relevant exposure surfaces in SARS-CoV-2 BioSample metadata.

### 1.1 Related Work and Research Gaps

Research on genomic cybersecurity has identified integrity risks across sequencing workflows, analytical pipelines, and public genomic repositories. In our previous study [14], we examined cybersecurity risks across the complete next generation sequencing workflow and identified integrity weaknesses at several stages of data generation, processing, and analysis. Vinatzer et al. [15] explored the risks created by open submission processes and limited verification in pathogen genome databases, while Schumacher et al. [16] examined related integrity concerns in sequencing pipelines and public repositories. Caswell et al. [17] further argued that biological databases should be treated as global critical infrastructure because of their importance to scientific research, disease surveillance, and public health response. More closely related to the present study, Pauwels [13] described biological datasets and biotechnology infrastructure as potential targets of covert manipulation and adversarial information operations. Fouad [12] similarly examined cyberbiosecurity risks arising from the compromise and exploitation of biological information. Together, these studies show that biological information can become a strategic target rather than only a technical, privacy, or data quality concern. However, they do not provide a phase structured adversarial lifecycle, integrate manipulation across genomic data, analytical models, digital infrastructure, and governance systems, or evaluate these risks against metadata held in an operational genomic repository.

Metadata quality and data provenance present further integrity challenges. Gonçalves et al. [7] analysed 6.6 million BioSample records and identified substantial inconsistencies in submitted metadata. These inconsistencies can affect sample interpretation, record linkage, and downstream analysis. At the software level, Ney et al. [18, 19] showed that sequencing instruments and their associated software create additional attack surfaces within genomic workflows. Other studies have highlighted limitations in provenance tracking, workflow interoperability, and genomic data sharing infrastructures [20–22]. Collectively, this literature shows that genomic data integrity depends not only on sequence accuracy, but also on reliable metadata, software, provenance records, and data exchange processes. However, these weaknesses have not been examined together through an adversarial framework or assessed empirically at repository scale.

A related body of research has examined the growing role of artificial intelligence in biosecurity. Zhang et al. [23] demonstrated that DNA language models can be manipulated through jailbreak attacks to generate sequences with pathogenic characteristics. Killoran et al. [24] showed that deep generative models, including generative adversarial networks, can produce synthetic DNA sequences that preserve structural features of natural sequences and can be directed towards selected properties. Brent and McKelvey [25] argued that foundation models are lowering the level of technical expertise required to generate biologically relevant sequences. Similarly, the National Institute of Standards and Technology (NIST) [26] noted that advances in AI-driven nucleic acid design are developing faster than some existing biosafety safeguards. Policy assessments have also highlighted the growing risk of biological misuse associated with frontier AI systems [27]. Ranjan et al. [28] examined the broader biocybersecurity landscape, while Walsh [29] discussed privacy and security risks in genomic data-sharing systems. Collectively, this literature shows that generative AI has increased the accessibility and scale of biological sequence generation, creating new challenges for screening, verification, and genomic data integrity. However, limited attention has been given to how AI-generated or manipulated data could be introduced gradually into open genomic repositories as part of a persistent and coordinated threat.

Research on adversarial ML shows that analytical systems can be degraded through corrupted, manipulated, or strategically biased data. Alber et al. [9] found that poisoning a small proportion of a medical language model’s training data increased harmful outputs. Annan et al. [30] identified related security risks in genomic ML systems that depend on large biological datasets and complex analytical workflows. Finlayson et al. [31] demonstrated that adversarial manipulation can alter medical ML predictions, while Schwarzschild et al. [32] benchmarked backdoor and data poisoning attacks under consistent conditions and showed that their effectiveness depends heavily on the training setup. Although these studies examine different analytical settings, they consistently demonstrate the sensitivity of ML systems to manipulated data.

Research on backdoor attacks, model inversion, distributed learning vulnerabilities, and concept drift further shows that ML systems face security and reliability risks throughout model development and deployment [33–39]. Reconstruction and model inversion are not data poisoning attacks, but they illustrate the broader range of vulnerabilities affecting ML systems. More directly, Koilakos et al. [40] examined targeted poisoning and backdoor attacks against genomic foundation models, including sequence poisoning during pretraining and label corruption in BRCA1 variant classification. Their work provides empirical evidence that genomic ML models are vulnerable to targeted poisoning. However, it focuses on genomic foundation models and clinical variant interpretation rather than pathogen surveillance, repository metadata, or persistent manipulation across multiple layers of a surveillance system.

Overall, the literature shows that genomic repositories contain integrity and provenance weaknesses, generative AI has expanded access to synthetic sequence generation, biological information can be strategically targeted, and ML models are sensitive to poisoned data. However, these areas remain largely disconnected. Existing research does not provide a unified lifecycle for examining persistent manipulation across genomic repositories, metadata, analytical models, digital infrastructure, and governance, nor does it assess, at repository scale, whether the metadata conditions such a campaign would depend on are already present in pathogen surveillance data.

### 1.2 Contributions of this Paper

To address the identified gap, this paper combines a conceptual threat framework with a large-scale empirical audit of operational genomic repository metadata. The main contributions are as follows:

- **The APBT threat concept.** We introduce APBTs as a data-centric cyber-biological threat concept, in which an adversary degrades genomic surveillance by acting on sequences, metadata, reference datasets, analytical models, and governance systems rather than on biological material itself. While prior work has examined repository integrity, synthetic sequence generation, and adversarial ML as separate technical problems [7, 15–17, 23, 24], to our knowledge none has framed them as components of a single, strategically coordinated campaign against genomic surveillance.
- **Development of an integrated APBT lifecycle and taxonomy:** The paper proposes an integrated multilayer APBT taxonomy organised around a four-phase adversarial lifecycle: strategic planning and reconnaissance, initial infiltration and positioning, persistent poisoning, and activation or operational disruption. The lifecycle adapts the planning logic of APTs in cybersecurity [10, 11] to the informational infrastructure of public health, providing a structured basis for analysing threats that develop gradually rather than as discrete incidents.
- **A systematic metadata audit framework.** We conduct an empirical assessment of the metadata component of the APBT framework using 23 checks across five metadata layers: temporal, geographic, host and specimen, provenance and administrative, and technical and analytical metadata. The checks are informed by community metadata standards [41, 42]. This provides a reproducible method for identifying metadata conditions that reduce record verification, provenance reconstruction, and anomaly detection.
- **Repository-scale empirical evidence of metadata exposure.** In contrast to Gonçalves et al. [7], who examined metadata quality mainly in relation to standardisation, data discovery, and reuse, we provide an adversarial security assessment of 100,000 SARS-CoV-2 BioSample records. By applying the proposed audit framework, we demonstrate how metadata incompleteness can act as an APBT-relevant exposure surface that weakens verification, provenance reconstruction, traceability, and the detection of manipulated records, particularly within the provenance and technical metadata layers.

### 1.3 Paper Organization

The paper is organised as follows. Section 2 presents the background and APBT lifecycle framework; Section 3 describes the metadata vulnerability audit of 100,000 SARS-CoV-2 BioSample records and discusses the implications of the findings; and Section 4 concludes the paper and highlights future work directions.

## 2 Background and APBT Lifecycle Framework

This section develops the conceptual foundation of the APBT framework. It first explains how APBTs extend the logic of APTs from conventional cybersecurity into genomic surveillance and cyberbiosecurity, where the primary target is not digital infrastructure alone but the informational systems that support public health interpretation. Table 1 summarises the key distinctions between traditional APTs and APBTs across ten analytical dimensions. It then introduces the APBT lifecycle and multilayer taxonomy, showing how adversarial activity can progress from strategic planning and initial positioning to persistent poisoning and operational disruption. This conceptual framework provides the basis for the empirical metadata audit presented in the following section.

**Table 1:** Comparison between traditional APTs and APBTs across ten analytical dimensions.

| Dimension | Traditional APT | APBT |
| --- | --- | --- |
| **Primary target** | Computational and organisational infrastructure, including networks, servers, endpoints, software, credentials, and internal data systems. | Genomic surveillance information systems, including sequences, metadata, reference datasets, lineage labels, analytical models, and repository governance processes. |
| **Entry point** | Phishing, zero-day exploits, malware, stolen credentials, or direct compromise of digital infrastructure. | Open repository submissions, metadata fields, reference updates, annotation changes, model training datasets, or shared bioinformatics workflows. |
| **Main object of manipulation** | Code, configurations, credentials, firmware, system logs, network traffic, or access permissions. | Genomic sequences, metadata, labels, lineage definitions, reference datasets, model inputs, or analytical outputs. |
| **Skills required** | Cyber intrusion, exploit development, persistence, privilege escalation, and evasion skills. | Bioinformatics expertise, genomic data manipulation, metadata design, ML knowledge, and the ability to generate scientifically plausible records. |
| **Persistence strategy** | Maintain covert access inside target systems over an extended period. | Gradually introduce plausible poisoned data, misleading metadata, or biased analytical inputs that persist through normal repository and pipeline workflows. |
| **Detection challenge** | Can be investigated through intrusion detection, log analysis, endpoint monitoring, behavioural anomaly detection, and network security controls. | Harder to detect using conventional cyber tools because manipulated records may appear scientifically plausible and may pass routine metadata or format validation. |
| **Operational objective** | Data theft, espionage, sabotage, disruption of IT systems, or degradation of organisational capability. | Distortion of biosurveillance outputs, delayed outbreak detection, weakened variant classification, incorrect provenance inference, and reduced confidence in public health decisions. |
| **Consequences** | Loss of confidentiality, integrity, or availability of digital systems and organisational data. | Distorted biological interpretation, flawed phylogenetic inference, biased epidemiological modelling, inaccurate diagnostic or surveillance outputs, and weakened preparedness. |
| **Role of AI** | Automation of reconnaissance, phishing, vulnerability discovery, malware generation, evasion, or intrusion support. | Generation of plausible synthetic sequences, metadata patterns, adversarial labels, poisoned training examples, or misleading analytical outputs [23, 25, 43]. |
| **Barrier to entry** | Often high because long-term covert access usually requires advanced technical capability and operational infrastructure. | Potentially lower for repository-based attacks because open data ecosystems, public tools, and generative AI can be exploited without wet-lab access. |

### 2.1 From APTs to APBTs

APBTs represent an emerging class of risk at the intersection of cybersecurity, biological science, and public health. An APBT is a long-duration, data-centric, cyber-biological campaign that incrementally poisons genomic information systems to erode trust in public health decision making, distort surveillance outputs, and conceal adversarial activity without direct biological manipulation. APBTs can be conceptualised as the biological surveillance analogue of APTs in cybersecurity.

Traditional APTs are long-duration, strategically motivated cyber campaigns in which adversaries maintain covert access to computational systems and use incremental actions to achieve objectives that remain concealed until a later point of advantage [10, 11]. APBTs follow a similar strategic pattern, but their target is different. Whereas APTs focus on networks, servers, software, credentials, and organisational data systems, APBTs target the informational infrastructure of genomic surveillance: genomic sequences, metadata, reference datasets, analytical models, lineage assignments, and governance processes. The threat therefore acts through data and interpretation rather than through direct alteration of biological materials. APBTs do not require unauthorised system penetration or laboratory-level manipulation. An adversary can instead introduce misleading sequences, corrupted metadata, biased reference updates, or subtle analytical distortions into genomic repositories and shared bioinformatics resources. Each contribution can appear scientifically plausible when viewed in isolation, allowing manipulation to remain difficult to distinguish from ordinary data variation, incomplete metadata, contamination, or routine submission error. The critical difference between an APBT and innocent error lies in strategic coordination over time: poisoned data are introduced according to a concealed objective, with patterns designed to resist normal curation and produce a specific bias at a future point of operational importance.

The strategic concern is that APBT activity can begin long before a biological event or public health emergency becomes visible. By gradually poisoning genomic datasets, metadata fields, reference files, and analytical models, an adversary weakens the systems responsible for detecting abnormal genetic patterns, classifying variants, and identifying unusual sequence features. The COVID-19 pandemic illustrates why this matters. The rapid sequencing and open sharing of the SARS-CoV-2 genome supported early diagnostic assay design, vaccine development, and global variant surveillance [1–3]. If the repositories, reference datasets, or metadata systems on which these analyses depended had already been systematically weakened or contaminated, early interpretation of the virus could have been delayed, variant classification could have been less reliable, and public health responses could have been based on incomplete or misleading genomic evidence. When a genuine outbreak occurs, a compromised surveillance system can misclassify variants, misinterpret evolutionary signals, or fail to generate timely warnings. This creates confusion and delay at a moment when public health responses depend on accurate genomic interpretation. In essence, an APBT is a cyber-biological threat that weaponises data rather than pathogens.

This comparison establishes APBTs as a distinct form of persistent threat directed at the informational foundations of genomic surveillance. The next section develops this concept further by presenting the APBT lifecycle and showing how adversarial activity progresses from planning and positioning to persistent poisoning and operational disruption.

### 2.2 The APBT Lifecycle and Multilayer Threat Taxonomy

The APBT lifecycle describes how a persistent cyber-biological campaign develops over time, from early strategic planning to operational disruption during a public health crisis. The lifecycle is organised into four phases: strategic planning and reconnaissance, initial infiltration and positioning, persistent poisoning, and activation and operational disruption. The overall lifecycle and taxonomy are illustrated in Figure 1.

**Figure 1:**
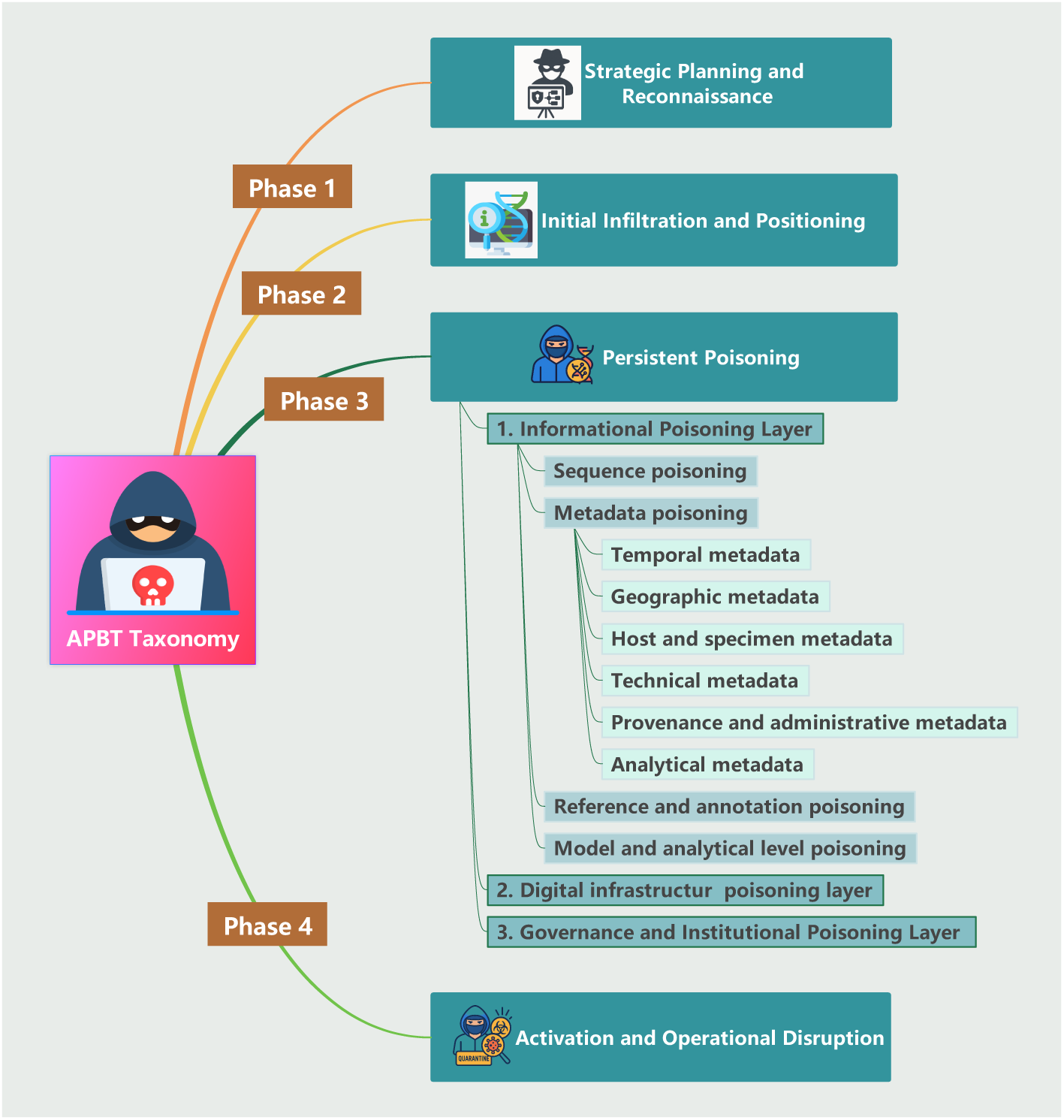
APBT taxonomy showing the four-phase lifecycle and the main poisoning layers involved in persistent threats to genomic surveillance.

#### 2.2.1 Phase 1: Strategic Planning and Reconnaissance

The first phase involves long-term analysis of the genomic surveillance ecosystem and the definition of the campaign’s strategic objective. In simple words, an APBT begins before any data are manipulated. The adversary first studies the genomic surveillance ecosystem, identifies its most influential components, and defines the long-term objective of the campaign. This reflects the planning logic of traditional APTs, in which attackers observe the target environment and locate weaknesses before their broader purpose becomes apparent [10, 11]. In an APBT, the objective may be broad, such as gradually weakening trust in a widely used genomic resource, or more focused, such as distorting the interpretation of a particular pathogen, lineage, or geographic signal at a strategically important time.

The adversary is therefore concerned less with gaining network access and more with understanding how information moves through the surveillance system. This includes identifying which repositories are most trusted, which metadata fields influence public health interpretation, how reference datasets are updated, which analytical pipelines are widely reused, and how scientific outputs inform surveillance dashboards and policy decisions. High-value targets are those that combine influence with limited verification, such as an INSDC member repository, a high-volume submitting institution, a widely adopted lineage-definition resource, or a pipeline whose outputs are reused across multiple surveillance systems. Manipulation at any one of these points can spread through downstream analyses and affect a much wider part of the ecosystem. By the end of this phase, the adversary has defined the campaign objective, selected the target, and determined the form of distortion to be introduced in later stages.

#### 2.2.2 Phase 2: Initial Infiltration and Positioning

Following the strategic planning stage, the APBT lifecycle progresses to initial infiltration and positioning, where the adversary quietly enters the targeted genomic information systems and becomes part of routine data flows. The purpose of this phase is not to manipulate information immediately, but to establish a low-visibility foothold that could enable sustained activity in later stages. By interacting with the system in ordinary, scientifically plausible ways, the adversary gains familiarity with operational norms while avoiding the scrutiny that accompanies anomalous behaviour. Because these ecosystems are designed for openness and rapid data sharing, initial participation often faces minimal barriers. The adversary leverages this openness to integrate smoothly into existing processes, ensuring future submissions appear consistent with expected patterns. To remain unnoticed, the adversary engages in activities that resemble legitimate use, such as submitting routine or benign data. Over time, these interactions may create a behavioural baseline against which later submissions appear normal. This gradual normalisation is essential: it reduces the likelihood that subsequent manipulations will stand out or trigger additional scrutiny from automated validation systems or human curators. Section 3 demonstrates why this normalisation is feasible in practice. Because most records do not identify the project or institution from which they originate, incomplete and low-context submissions are the norm across the repository. A new submitter therefore, does not appear unusual, as there is no established provenance baseline against which anomalous participation could be identified.

Initial infiltration also allows the adversary to study how the ecosystem behaves under real operating conditions. Unlike the external reconnaissance of Phase 1, the adversary can now observe these processes from within: by monitoring how submissions are validated, how metadata is handled, and how quickly information propagates across pipelines and repositories, the attacker develops an understanding of system tolerances and workflow structures. This insight is crucial for identifying where manipulations can later be introduced with minimal risk of detection. This stage therefore establishes the practical foundation needed for the long-horizon poisoning activities characteristic of APBT.

#### 2.2.3 Phase 3: Persistent Poisoning

Phase 3 forms the operational core of an APBT campaign. At this stage, the adversary begins to introduce gradual and coordinated distortions into the systems that support genomic surveillance and public health decision making. The aim is not to cause an immediate or easily recognised failure, but to erode data integrity, analytical reliability, and institutional trust over time, often long before an outbreak or other biological event becomes visible. Persistent poisoning operates across three interconnected layers: (i) informational poisoning, (ii) digital infrastructure poisoning, and (iii) social, institutional, and governance poisoning.

##### Informational Poisoning

The informational layer forms the foundation of an APBT because it targets the data on which all genomic analysis relies. At this layer, the adversary manipulates the scientific inputs used for surveillance and interpretation, including genomic sequences, metadata, reference datasets, annotation files, and the training data used by ML models. Altering these inputs, even subtly, can influence downstream analyses and decision making without any interference with laboratory operations or computational infrastructure. This informational poisoning operates through four distinct but interconnected mechanisms: (i) sequence poisoning, (ii) metadata poisoning, (iii) reference and annotation poisoning, and (iv) model and analytic poisoning, which are examined in turn below.

###### I. Sequence poisoning

Sequence-level poisoning involves the intentional introduction of manipulated, fabricated, or misleading genomic sequences^1^ into public or shared datasets. Because automated quality control validates format rather than biological plausibility or provenance, poisoned sequences that appear technically valid can pass initial checks [14, 15, 17]. An adversary therefore requires no access to biological materials, exploiting instead the openness of submission systems and the automation that governs data ingestion. Once accepted, poisoned sequences propagate through phylogenetic workflows, reference updates, ML training cycles, and public health surveillance dashboards, amplifying their impact over time. These characteristics make sequence-level poisoning a foundational component of an APBT, capable of weakening global genomic surveillance long before a biological event is detected.

It can take several forms: introduction of fully synthetic sequences generated by sequence language models such as GenSLM and HyenaDNA or by GAN based systems, which produce natural looking genomic regions; engineered variant insertion, embedding artificial mutations that mimic natural change; contaminant or paralog substitution, submitting plausible sequences that do not originate from the claimed organism; random noise or low complexity insertion, which quietly shifts analytical baselines; phylogenetic misplacement, constructing sequences that distort evolutionary trees and imply false transmission links; and stealth alteration of widely used reference sequences, whose effects become ecosystem wide once incorporated into reference releases. In the COVID-19 context, even a few falsified early SARS-CoV-2 genomes could have introduced artificial branches into phylogenetic trees or obscured the detection of a genuinely emerging variant.

###### II. Metadata poisoning

Every sequencing experiment produces two components: the biological sequence itself (stored in formats such as FASTQ, BAM, or FASTA) and the metadata describing sample origin, handling, and analysis [45]. The sequence carries the biological signal, whereas the metadata supplies the context needed to interpret, compare, and trace it. Metadata poisoning is the deliberate manipulation of these descriptive fields. Because analysis pipelines, epidemiological models, and ML systems depend on metadata for contextual interpretation, even small distortions can cascade into large analytical and policy errors [46, 47]. This risk is heightened because most repositories accept fields such as dates and locations at face value, rarely verifying them against laboratory records. The following subsections describe how poisoning manifests across individual metadata fields, and Table 2 summarises the categories and associated attack vectors.

***a. Temporal metadata***, including collection date, sequencing date, and submission date, defines the timeline used for mutation-rate estimation, phylogenetic reconstruction, and surveillance trend analysis. If these dates are inaccurate or deliberately altered, a lineage can appear older or newer than it is, or seem to arrive in a region at the wrong time, biasing molecular-clock models and misrepresenting when a variant emerged, how quickly it spread, and whether a signal reflects contamination, introduction, or genuine evolution. These distortions are especially consequential during public health emergencies such as COVID-19, when variant detection and transmission assessment depend on accurate sampling timelines. Because date fields are rarely verified against laboratory logs or instrument records, falsified timestamps can enter repositories and influence downstream analyses without triggering immediate warnings.
***b. Geographic metadata*** records where a biological sample was collected, including fields such as country, region, and collection site. This spatial information supports routine research on regional genetic diversity, microbial ecology, and evolutionary patterns, and it is essential during outbreaks for tracing transmission, identifying where new variants emerge, and directing public health measures such as testing and travel guidance [46, 47]. When geographic metadata is poisoned, both scientific inference and emergency response can be disrupted. A single mislabelled sample can create the false impression that a pathogen has appeared in a new region or hide activity in the true location. When multiple manipulated entries accumulate, they can generate artificial transmission routes, obscure genuine outbreak clusters, and distort phylogeographic analyses. An adversary could exploit this by fabricating international spread, influencing travel restrictions, creating political tension, or masking an emerging hotspot. Even small shifts in location can bias research findings or degrade ML models that incorporate spatial features. Detection is difficult because repositories and pipelines generally accept geographic fields without independent verification.
***c. Host and specimen metadata*** specifies the biological source of a sequence, indicating whether it came from a human, animal, or environmental sample and identifying the specimen type. These fields are essential for interpreting pathogen ecology, transmission pathways, and possible zoonotic events. During COVID-19, host and specimen information helped distinguish human cases from animal or environmental samples, enabling reconstruction of potential spillover routes and assessment of environmental persistence. Manipulating these fields, for example by relabelling an animal or environmental sample as human, or a laboratory isolate as a clinical specimen, could have misdirected investigations into the origin and spread of SARS-CoV-2, distorting reconstruction of spillover routes or masking laboratory-associated sequences.
***d. Technical metadata*** captures the operational characteristics of a sequencing run, including read counts, read length, quality scores, flow-cell identifiers, and platform information. These fields are widely used for automated quality control, instrument comparison, and ML-based filtering. Because modern sequencing platforms and laboratory information systems generate and upload these fields automatically, they often receive limited manual scrutiny. An adversary can exploit this automation by altering technical metadata while keeping values numerically plausible, for example inflating read counts to conceal low-quality data, assigning misleading platform identifiers, or generating combinations of metrics that appear valid but introduce subtle inconsistencies. Such manipulations distort quality assessments, degrade benchmarking studies, and corrupt downstream analytical models, and they are difficult to detect because the altered values frequently fall within normal statistical ranges.
***e. Provenance and administrative metadata*** describes the institutional, procedural, and submission context in which genomic data is generated and shared [48]. Manipulating provenance fields can have serious consequences. An adversary could submit data under a falsified or impersonated laboratory name, alter project identifiers to obscure the true origin of a dataset, or attach data to the BioProject of a respected institution to create false legitimacy. If a submission account is compromised, poisoned data may enter repositories through an apparently trustworthy channel. Even small inconsistencies in laboratory names, email domains, or project codes can confuse curators and complicate later auditing. The scale of this exposure is documented in Section 3, where institutional attribution collected_by was absent from 51.3% of records and project-level BioProject linkage from 99.3%, leaving little provenance context against which such manipulation could be checked.
***f. Analytical metadata***, including software versions, reference identifiers, and pipeline parameters, governs reproducibility. Misreporting a version or altering a reference identifier leaves the sequence untouched but changes how it is interpreted, causing systematic miscalls or lineage misassignment. For instance, silently substituting the reference genome version used for lineage assignment could cause a genuine variant to be misclassified across every record processed with that pipeline, while the underlying sequences remain entirely valid. Effective mitigation requires cross-field consistency checks, validation against instrument-generated and pipeline execution records, and provenance mechanisms that ensure analytical fields accurately reflect how each sequence was processed.

**Table 2:** Threat model: metadata categories and associated poisoning risks, with illustrative COVID-19 examples.

| Metadata category | Poisoning vector | Impact on analysis and public health | Illustrative COVID-19 example |
| --- | --- | --- | --- |
| **Temporal** | Manipulated collection, sequencing, or submission dates. | Biased molecular-clock estimates, distorted variant emergence timelines, and flawed transmission trends. | Shifting collection dates to make a variant appear to emerge earlier than it did. |
| **Geographic** | False or masked geographic origin. | Distorted transmission maps, artificial transmission routes, and flawed phylogeographic models. | Changing sample location to imply spread into a new region. |
| **Host and specimen** | False host information, incorrect specimen source, or fabricated lineage. | Incorrect evolutionary interpretation, false host associations, and misdirected spillover investigation. | Relabelling a laboratory or animal sample as a human clinical specimen. |
| **Technical** | Altered sequencing platform, reagent batch, or quality metrics. | Misleading error profiles, incorrect variant calls, and biased quality control. | Falsifying coverage so that missing mutations go undetected. |
| **Provenance and administrative** | Falsified submitter identity, laboratory name, or project identifiers. | Breakdown of provenance, reduced traceability, and loss of accountability. | Uploading engineered sequences under a legitimate laboratory identity. |
| **Analytical** | Manipulated software versions, falsified reference identifiers, or altered parameters. | Systematic variant miscalling and incorrect lineage assignment. | Changing the reference genome version used for lineage classification. |

###### III. Reference and annotation poisoning

Reference datasets, including reference genomes such as GRCh38.p14 [49], curated mutation lists, and lineage definitions such as those maintained by PANGO [50], provide the coordinate system for alignment and the basis for lineage assignment. Poisoning them disrupts analysis ecosystem-wide: manipulated consensus genomes shift alignment coordinates, altered lineage definitions cause systematic misclassification, and corrupted taxonomies undermine automated identification. These risks are amplified because reference datasets are reused widely and updated automatically, often without independent verification. A poisoned SARS-CoV-2 lineage definition, for instance, could have caused a genuine variant of concern to be silently misassigned across every pipeline that adopted the update.

###### IV. Model and analytic poisoning

Model and analytic level poisoning targets the computational systems that convert genomic data into surveillance outputs. Instead of breaking a model directly, the adversary manipulates the data, labels, parameters, or input distributions on which the model depends, causing outputs to become gradually biased or unreliable [9, 31, 40]. In genomic surveillance, this can affect lineage classifiers, quality filters, anomaly detectors, and outbreak forecasting tools. The main risk comes from poisoned training data. If corrupted sequences, incorrect labels, or misleading metadata enter public datasets, models can learn false relationships between genomic patterns and surveillance categories. A lineage classifier repeatedly exposed to incorrect labels, for example, may slowly shift its decision boundary while still appearing accurate on routine validation data. Similar risks occur in federated learning, where institutions collaboratively train models by sharing updates rather than raw data [51]. In adversarial settings, a malicious participant can bias the shared model through manipulated updates while preserving apparently normal performance [52]. More covert mechanisms include backdoor poisoning, where hidden patterns cause a model to behave normally on standard inputs but incorrectly when a specific trigger condition is present [33, 34], and concept drift exploitation, where small repeated changes in the data distribution gradually redefine what the model treats as normal [38, 39]. Model inversion and data leakage are mainly privacy attacks, but they can also support poisoning cycles if reconstructed training-like examples are modified and reintroduced into future datasets [36, 37]. In a COVID-19-like outbreak, this type of poisoning could cause a surveillance model to treat an important variant signal as ordinary background variation, misclassify a lineage, or delay warning generation. The result is not an obvious model failure, but a gradual loss of sensitivity when accurate genomic interpretation is most needed.

##### Digital Infrastructure Poisoning Layer

Poisoned genomic information has limited impact if it remains confined to a single record or system. Its influence grows when the digital infrastructure of genomic surveillance processes, copies, distributes, and repeatedly reuses it. This layer therefore concerns the systems through which genomic information moves, including bioinformatics pipelines, cloud platforms, automated learning systems, surveillance dashboards, and distributed repositories. Unlike informational poisoning, which directly targets sequences, metadata, reference datasets, or analytical inputs, digital infrastructure poisoning affects the environments that transform and circulate these resources. Manipulation at this layer can allow a small distortion to persist across workflows, spread between institutions, and influence downstream surveillance without altering biological material.

Five mechanisms are particularly important:

i. **Bioinformatics pipeline poisoning.** Bioinformatics pipelines convert raw sequencing signals into assembled genomes, variant calls, lineage assignments, and other surveillance outputs. Because these workflows rely on interconnected software tools, instrument components, configuration files, and external dependencies, a small change at one stage can influence every result produced downstream. Manipulation of base-calling parameters, filtering thresholds, reference-matching rules, workflow definitions, or trusted software packages can alter analytical behaviour while leaving the pipeline apparently functional [14, 18]. The resulting errors may resemble ordinary instrument variation or software behaviour, making persistent manipulation difficult to recognise.
ii. **Cloud infrastructure poisoning.** Cloud platforms increasingly serve as shared environments for storing genomic data, running analytical workflows, and distributing reference resources. This centralisation improves access and scalability, but it also concentrates risk. Compromised credentials, excessive permissions, or altered cloud-hosted files can introduce corrupted sequences, metadata, or reference datasets into several workflows at once [53, 54]. Once a manipulated resource is treated as authoritative, every laboratory or pipeline that reuses it can inherit the same distortion.
iii. **Automated retraining poisoning.** Genomic classifiers, lineage detectors, and forecasting systems are increasingly updated as new sequences and metadata become available. This continuous adaptation becomes a weakness when manipulated records enter automated ingestion and retraining cycles. Rather than attacking the model directly, the adversary alters the data environment surrounding it. Repeated exposure to poisoned inputs can gradually shift decision boundaries, redefine statistical baselines, and make abnormal genomic patterns appear routine. Automated retraining therefore acts as a bridge between informational poisoning and model-level degradation, allowing analytical systems to internalise and amplify corrupted data through their own updating processes [55].
iv. **Dashboard manipulation.** Surveillance dashboards are where complex genomic analyses are translated into signals that researchers, public health agencies, and policy makers can act upon. By combining repository data, analytical outputs, and epidemiological indicators, they provide a concise view of variant prevalence, mutation patterns, and geographic spread. This visibility also makes them a critical point of vulnerability. Manipulation of data feeds, back-end databases, analytical summaries, display thresholds, or administrative settings can change the picture presented to users [55, 56]. As a result, the underlying data may remain partly accurate while the operational view becomes misleading. Since many decisions are based on dashboard outputs rather than raw genomic records, distortion at this interface can directly influence interpretation, prioritisation, and public health response.
v. **Mirror repository poisoning.** Global genomic data sharing depends on distributed repositories and institutional mirrors that improve availability and reduce access delays [57]. However, mirrored systems may differ in synchronisation frequency, security controls, provenance validation, and integrity checking. These differences can allow an altered or outdated copy to diverge from the authoritative source while continuing to appear legitimate. Researchers who retrieve data from the affected mirror may then incorporate localised distortions into their analyses without recognising that the source has changed [7]. Mirror poisoning is therefore especially difficult to diagnose because the primary repository may remain intact while analytical errors emerge only in particular institutions or regions.

Digital infrastructure poisoning acts as the propagation and amplification layer of an APBT. Informational poisoning introduces the distortion, but infrastructure determines how widely it travels, how long it persists, and how often it is reproduced. By exploiting interconnected pipelines, cloud services, learning systems, dashboards, and repository mirrors, an adversary can transform a limited manipulation into a broader surveillance failure. The significance of this layer lies not only in the compromise of individual systems, but in the capacity of trusted infrastructure to repeatedly process and distribute poisoned information across the genomic surveillance ecosystem.

##### Social, Institutional, and Governance Poisoning Layer

The governance layer represents the highest level of manipulation within an APBT. Rather than targeting data or computation, it targets the institutional mechanisms that maintain trust, accountability, and coordination, and attacks here are structurally damaging because they erode the very safeguards that would otherwise catch poisoned submissions. The main vectors are: *credential compromise and account misuse*, where phishing or password reuse lets an attacker upload falsified data or impersonate a trusted institution through a legitimate channel; *provenance tampering*, altering collection dates, project identifiers, or laboratory names that are often stored without cryptographic signing; *manipulation of logs and audit trails*, deleting or fabricating the version histories curators rely on to reconstruct how data were altered; *identity and authorship spoofing*, submitting under the names of trusted institutions to exploit reliance on reputation; *supply chain manipulation of governance tools*, compromising laboratory information systems or metadata templates so that all future submissions carry falsified provenance with minimal ongoing effort; and *Sybil and impersonation attacks*, creating many fraudulent accounts to submit poisoned data at scale or spoofing trusted institutions to lend it credibility.

#### 2.2.4 Phase 4: Activation and Operational Disruption

Phase 4 is the stage at which accumulated poisoning is activated, not by any new action from the adversary, but by the operational pressure of a real event such as a COVID-19-like outbreak or the emergence of a novel pathogen. The dormant corruption seeded across earlier phases now becomes operative: the volume and stakes of incoming data rise sharply, and a compromised analytical baseline that had lain inert must interpret a flood of genuinely novel signals at speed. The resulting damage has two distinct sources that compound each other. First, the historical baseline is already biased, so new sequences are aligned, classified, and dated against corrupted reference points, and legitimately anomalous patterns are read as background variation. Second, an adversary who remains active continues to introduce fresh distortions in real time, exploiting footholds established across pipelines, dashboards, and retraining workflows during earlier phases. The first source misreads good data; the second injects bad data; together they leave decision makers with inconsistent signals from different pipelines at precisely the moment accurate analysis matters most. Had such corruption been seeded through the early pandemic, the surveillance system would have entered the emergence of a variant such as Omicron already misaligned, interpreting a genuinely novel lineage through a compromised baseline.

Recovery is especially challenging because the damage is closely linked to the structural weaknesses identified in the empirical audit. Once metadata has been altered or contaminated, correcting it becomes extremely difficult without strong provenance records. As reported in Section 3, these links were missing from almost all of the audited BioSample records. Without a clear record of where the data came from, who submitted it, and how it was processed, there may be no reliable way to trace the problem, restore the original information, or rebuild an accurate historical record.

## 3 Large-Scale Metadata Vulnerability Audit of SARS-CoV-2 BioSample Records

To assess whether the metadata weaknesses described in the APBT taxonomy exist in operational repositories, we audited 100,000 SARS-CoV-2 BioSample records from NCBI against 23 checks spanning five metadata layers. This section reports the completeness, provenance, and traceability of these records and evaluates the structural conditions relevant to APBT-style manipulation.

### 3.1 Audit Framework and Methodology

#### 3.1.1 Data Collection

To empirically validate the APBT vulnerability framework, a large-scale metadata audit was conducted on SARS-CoV-2 BioSample records from NCBI. NCBI BioSample constitutes one of the world’s primary repositories for biological sample metadata and is a core component of the INSDC, which also encompasses ENA and DDBJ [58, 59]. Records submitted to any INSDC member are mirrored across all three repositories, making NCBI BioSample a representative and operationally critical source for this analysis.

Data were retrieved programmatically using the NCBI Entrez API via Biopython’s Entrez module, following NCBI’s published usage guidelines [59]. The retrieval query targeted SARS-CoV-2 BioSample records using the canonical NCBI taxonomy identifier txid2697049[Organism]. This identifier-based approach was adopted over text-based searches because taxonomy IDs are permanent and immune to changes in organism name indexing, ensuring full reproducibility. All data were retrieved in read-only mode using publicly available interfaces, and no modifications were made to repository records.

A total of 100,000 BioSample records were retrieved in batches of 200 (consistent with NCBI API rate recommendations) using the efetch function with XML output format. For each record, the full BioSample XML document was parsed to extract all available metadata fields. Extraction was restricted to BioSample XML attribute fields to ensure the audit evaluated repository-level completeness rather than downstream annotations [45, 60]. These fields were mapped to five APBT metadata layers: temporal, geographic, host and specimen, provenance and administrative, and technical and analytical. The final dataset comprised 100,000 records with collection dates spanning 2019 to 2026.

#### 3.1.2 Sampling Design and Dataset Composition

Records were retrieved in the order returned by the Entrez query rather than by random sampling from the full result set, and the audited dataset is therefore best characterised as a registration ordered window of the repository. Registration timestamps in the retrieved set span 13 October 2021 to 15 April 2026, but the distribution is strongly weighted towards the end of that range, with 77,691 records (77.7%) registered during 2025 and a further 19,672 (19.7%) during 2026. Only 2,637 records (2.6%) were registered before 2025. The window includes material brokered through all three INSDC members, comprising 50,236 ENA accessions, 49,345 NCBI accessions and 419 DDBJ accessions, and spans 42 country level labels and 104 submitting handles. Because retrieval followed accession order, submission batches were generally captured intact rather than sampled record by record. The retrieved set has a data cutoff of 15 April 2026, which is the most recent registration and update timestamp present in the audited records.

#### 3.1.3 Analysis Strategy

The analysis was structured around the APBT metadata vulnerability taxonomy introduced in Section 2.2.3, which organises genomic metadata into five layers corresponding to distinct APBT attack surfaces. The audit framework was purpose-built for this study, since no standardised instrument exists for evaluating genomic metadata against an adversarial threat model. The fields examined, however, are not arbitrary: they correspond to the contextual attributes defined as minimally necessary by community metadata standards, including the Genomic Standards Consortium’s Minimum Information about any (x) Sequence (MIxS) specification [41] and, for SARS-CoV-2 specifically, the PHA4GE contextual data specification for open genomic epidemiology [42], which identifies when and where a sample was collected as the two most critical attributes for genomic surveillance. The analytical importance of each field is likewise established in prior work, spanning sampling-date precision for molecular-clock inference [55], geographic completeness for outbreak reconstruction [61], and provenance and traceability in repository-integrity studies [7, 15, 17]. The framework’s novelty therefore lies not in identifying new fields but in reframing their completeness as a set of adversarial exposure surfaces and integrating them into a single, severity-scored, multilayer audit. Each of the five metadata layers, temporal, geographic, host and specimen, provenance and administrative, and technical and analytical, was operationalised as a set of concrete checks against the fields within it.

A systematic audit framework of 23 individual checks was developed and applied to every record in the dataset. These checks were grouped into the five layers: seven temporal checks (T1–T7), three geographic checks (G1–G3), five host and specimen checks (H1–H5), four provenance checks (P1–P4), and four technical and analytical checks (A1–A4). Each check was operationalised as a binary indicator per record (metadata vulnerability indicator present or absent). Composite vulnerability scores were calculated by summing the number of triggered checks (range 0–23), enabling quantitative comparison of vulnerability exposure across metadata layers, geographic regions, and submitting institutions. Records were classified into four severity tiers: Low (1–2), Medium (3–5), High (6–9), and Critical (*≥*10). Individual check criteria were set from documented biological and submission conventions where available, for example the accepted emergence date of SARS-CoV-2 for pre-emergence detection, and the severity thresholds were defined empirically to reflect the number of independent metadata layers a record fails simultaneously.

##### Temporal Checks (T1–T7)

Seven temporal checks were applied to evaluate the integrity and precision of collection date metadata. Check T1 identified records with missing or unparseable collection dates using flexible multi-format parsing supporting YYYY-MM-DD, YYYY-MM, and YYYY formats. Checks T2 and T3 flagged records with year-only and month-year precision respectively, both of which are insufficient for high-resolution molecular clock analysis and transmission modelling that require day-level sampling resolution when estimating evolutionary rates during outbreak-scale genomic surveillance [55, 62]. Check T4 identified records with collection dates pre-dating the accepted emergence of SARS-CoV-2 (before 1 November 2019). Check T5 flagged future-dated collections, and check T6 identified logically impossible cases where the collection date post-dated the submission date. Check T7 identified same-day collection and submission events, which represent unusually short metadata processing intervals that may indicate reduced provenance resolution or automated submission workflows. Together, checks T1–T7 capture missing temporal metadata, reduced timestamp precision, biologically implausible dates, and inconsistencies between collection and repository submission timelines that could distort outbreak reconstruction or molecular clock inference if exploited.

##### Geographic Checks (G1–G3)

Three geographic checks evaluated the completeness and internal consistency of location metadata. Check G1 identified records with missing geographic_location fields, including entries containing placeholder strings such as not provided, unknown, or not applicable. These placeholder values were treated as functionally missing because they provide no usable spatial provenance for downstream outbreak reconstruction. A standardised set of 22 placeholder strings was compiled through empirical inspection of repeated occurrence patterns observed during exploratory dataset profiling and documented BioSample submission conventions. Check G2 tested cross-field consistency between the geographic_location and country fields by extracting the country component from geographic_location using colon- or slash-delimited hierarchical location strings consistent with BioSample geographic-location conventions, and comparing this value with the dedicated country field. Check G3 separately identified placeholder-value metadata entries in the geographic_location field, distinguishing placeholder-based masking from genuinely blank fields to support APBT-specific analysis of metadata obfuscation patterns that may reduce geographic traceability. Together, checks G1–G3 capture missing spatial metadata, placeholder-based geographic masking, and inconsistencies between structured location fields that could distort phylogeographic inference or transmission pathway reconstruction if exploited [55, 61, 63].

##### Host and Specimen Checks (H1–H5)

Five checks evaluated host and specimen metadata. Check H1 flagged records with missing host fields, treating placeholder entries such as not collected’ and not applicable’ as functionally missing. Check H2 screened populated host fields against a predefined list of non-human and laboratory-associated designations (for example VERO cell lines and animal hosts such as bat, mink, cat, and dog) using exact-string matching, so only exact matches were flagged and differently formatted variants were not captured. Check H3 flagged missing isolation-source fields, again treating blank and placeholder values as functionally missing; because these placeholders were captured under H3, Check H4, which specifically targeted placeholder-valued isolation sources, returned no additional records. Check H5 identified biologically implausible isolation sources for a human respiratory virus, such as library’, cultured virus’, cell culture’, and laboratory’, which indicate laboratory-derived rather than primary clinical sampling. Together, H1–H5 capture missing or non-human host attribution and absent, masked, or laboratory-derived specimen-source information that may affect interpretation of transmission pathways and zoonotic origin.

##### Provenance Checks (P1–P4)

Four provenance checks assessed the traceability and chain-of-custody integrity of records. Check P1 identified records lacking a collected_by field, which is the primary institutional field establishing who collected the sample and supports attribution of sampling responsibility and verification of institutional provenance. Check P2 identified records without a BioProject accession visible within the BioSample attribute-level metadata. Check P3 identified records without a linked GenBank accession number within the BioSample metadata, meaning the associated nucleotide sequence is not directly linked through the BioSample record metadata. Check P4 identified records without an SRA accession visible within the BioSample metadata, meaning raw sequencing data are not directly referenced within the BioSample record metadata [58, 60]. Together, P2–P4 capture three levels of upward provenance linkage: project, sequence, and raw data that are essential for forensic traceability within the BioSample metadata layer in the APBT Phase 2 detection framework [15, 17, 48, 64].

##### Technical and Analytical Checks (A1–A4)

Four technical checks evaluated the completeness of analytical metadata. Check A1 identified records lacking a declared sequencing platform, which is required to interpret sequencing error structure and ensure comparability across variant-calling workflows [65, 66]. Check A2 identified records without a Pango lineage assignment [50], the widely used SARS-CoV-2 lineage classification framework. Check A3 identified records without coverage depth, which is required to assess variant-calling confidence and assembly reliability [65]. Check A4 identified records without a declared assembly method [62, 65, 66]. Together, these checks capture missing analytical metadata that may reduce confidence in sequence interpretation, lineage assignment reliability, and downstream phylogenetic comparability across surveillance datasets.

##### Composite Vulnerability Scoring

A composite vulnerability score was computed for each record by summing the number of vulnerability indicators triggered across all 23 checks (range 0–23). Records were classified into four severity tiers: Low (score 1–2), Medium (3–5), High (6–9), and Critical (*≥*10). This scoring framework provides a per-record metadata vulnerability exposure profile that supports identification of the most vulnerable records and enables comparison across submitting institutions and geographic regions.

The severity thresholds were set empirically based on the number of checks per layer (maximum 7 temporal + 3 geographic + 5 host + 4 provenance + 4 technical = 23), with High and Critical thresholds calibrated to reflect records failing checks across multiple layers simultaneously. Because each indicator corresponds to a distinct APBT metadata exposure surface as defined in the taxonomy, the composite score provides an interpretable approximation of the cumulative metadata vulnerability footprint associated with each BioSample record.

### 3.2 Results

#### 3.2.1 Overall Vulnerability Profile

Of the 100,000 SARS-CoV-2 BioSample records audited, every record (100.0%) contained at least one metadata field meeting the criteria for an APBT vulnerability indicator, demonstrating the presence of repository-level metadata exposure surfaces consistent with the assumptions of the APBT framework. The composite vulnerability score had a mean of 7.34 (SD 1.34) and a median of 8.0 out of a maximum possible score of 23, indicating that the average record triggered more than seven metadata vulnerability indicators. Severity classification showed that 95.9% of records fell within the High category (score 6–9) and 1.4% within the Critical category (score *≥*10), yielding a combined High and Critical rate of 97.3%. Only 2.7% of records were classified as Medium (score 3–5), and no records fell within the Low (score 1–2) or None (score 0) categories. The overall distribution of vulnerability severity and layer-level exposure rates is illustrated in Figure 2, and the results are summarised in Table 3.

**Figure 2:**
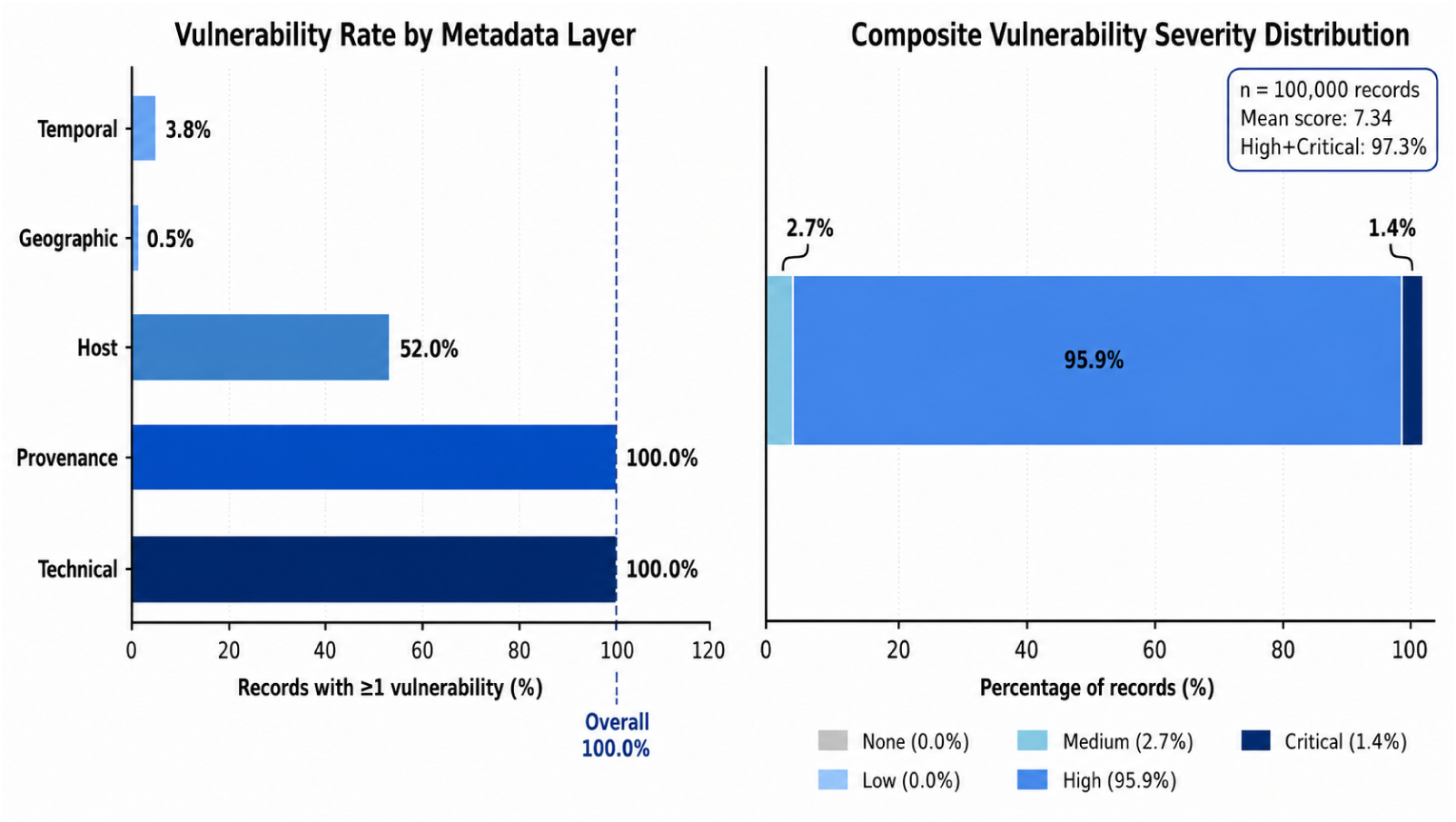
Vulnerability rate by APBT metadata layer (left) and composite severity distribution (right) across all 100,000 records. The provenance and technical layers reach 100% vulnerability, and 97.3% of records are classified High or Critical.

**Table 3:** Overall vulnerability summary statistics for 100,000 SARS-CoV-2 BioSample records from NCBI. Severity thresholds: Low = 1–2 failed checks; Medium = 3–5; High = 6–9; Critical *≥* 10.

| Metric | Value | % |
| --- | --- | --- |
| **Record counts** |  |  |
| Total records audited | 100,000 | 100.0 |
| Records with $\geq 1$ vulnerability | 100,000 | 100.0 |
| Records with no vulnerability | 0 | 0.0 |
| **Severity classification** |  |  |
| None (score = 0) | 0 | 0.0 |
| Low (score 1–2) | 0 | 0.0 |
| Medium (score 3–5) | 2,672 | 2.7 |
| High (score 6–9) | 95,922 | 95.9 |
| Critical (score $\geq 10$) | 1,406 | 1.4 |
| **Score statistics** |  |  |
| Mean vulnerability score | 7.34 | — |
| Median vulnerability score | 8.0 | — |
| Maximum vulnerability score | 12 | — |
| Total audit checks performed | 23 | — |

*Note.* Vulnerability score = number of indicators triggered out of 23 checks.

#### 3.2.2 Layer-by-Layer Vulnerability Findings

Vulnerability was not distributed uniformly across the five metadata layers. Table 4 summarises the rate at which each layer was flagged, the number of checks it comprises, and its correspondence to the APBT taxonomy; the following subsections examine each layer in turn. The provenance and technical layers were flagged in every record, whereas the temporal and geographic layers were flagged in fewer than 4% and 1% of records respectively, a pattern examined in detail below.

**Table 4:** Vulnerability rates by APBT metadata layer across 100,000 SARS-CoV-2 BioSample records. Each layer corresponds to an APBT poisoning attack surface as defined in the taxonomy.

| Layer | N flagged | % flagged | Checks | APBT taxonomy relevance |
| --- | --- | --- | --- | --- |
| **Temporal** | 3,804 | 3.8 | 7 | Phase 3, temporal poisoning: distorts molecular-clock estimates and variant emergence timelines. |
| **Geographic** | 479 | 0.5 | 3 | Phase 3, geographic poisoning: fabricates transmission routes and obscures genuine outbreak clusters. |
| **Host and specimen** | 51,963 | 52.0 | 5 | Phase 3, host and specimen poisoning: misrepresents zoonotic spillover and transmission pathways. |
| **Provenance and administrative** | 100,000 | 100.0 | 4 | Phase 2, infiltration: absence of provenance and chain-of-custody fields prevents detection of adversarial submissions. |
| **Technical** | 100,000 | 100.0 | 4 | Phase 3, analytical poisoning: prevents independent validation of sequencing results. |

##### Temporal Metadata Layer

The precision of collection-date metadata and the annual distribution of collection timelines are illustrated in Fig 3. Temporal metadata vulnerability indicators were identified in 3,804 records (3.8%). The majority of records provided full-precision collection dates (96,196 records; 96.2%), indicating strong temporal coverage across the dataset. However, 1,409 records (1.4%) provided only month–year precision (T3) and 1,279 records (1.3%) provided only year-level precision (T2), both of which are insufficient for high-resolution outbreak-scale molecular clock inference and transmission modelling [55, 67]. A further 874 records (0.9%) contained missing collection dates, and 242 records (0.2%) contained unparseable date strings.

**Figure 3:**
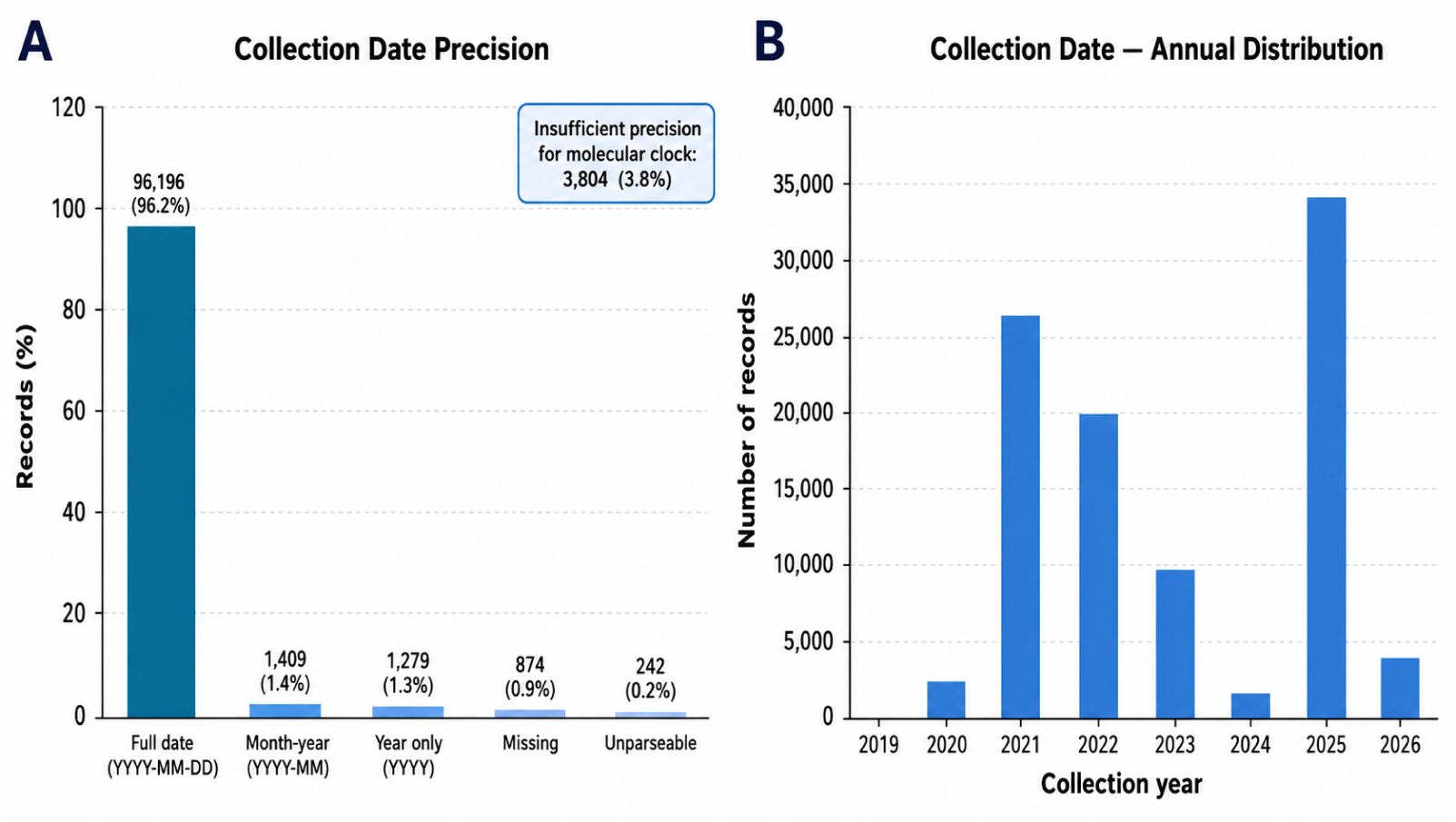
Temporal metadata analysis across 100,000 SARS-CoV-2 BioSample records. Left: collection-date precision, showing 96.2% full-precision dates (YYYY-MM-DD) and 3.8% (3,804) with insufficient precision for molecular-clock inference. Right: annual distribution of parseable collection dates (2019–2026), peaking in 2025, 2021, and 2022.

Importantly, no logically inconsistent temporal metadata were detected. No records contained pre-emergence collection dates (T4), future-dated collection timestamps (T5), or cases in which collection occurred after submission (T6). Likewise, no same-day collection and submission events (T7) were observed. These findings indicate that while reduced temporal precision represents a measurable metadata vulnerability surface, the dataset does not exhibit the extreme temporal inconsistencies typically associated with metadata entry errors or potential manipulation.

The annual distribution of collection dates spans 2019 to 2026, with peak submission volumes observed in 2021 (26,459 records), 2022 (19,997 records), and 2025 (34,322 records), reflecting sustained repository activity across multiple phases of the SARS-CoV-2 surveillance timeline.

##### Geographic Metadata Layer

Geographic metadata vulnerability indicators were the least prevalent across the five metadata layers, affecting 479 records (0.5%). These records contained placeholder values in the geographic_location field, predominantly not provided (381 records), together with not applicable (88 records) and Unknown (10 records). They were therefore flagged simultaneously by both G1 (missing location) and G3 (placeholder value), confirming that these checks identified the same subset of records. No inconsistencies between geographic_location and country fields (G2) were detected, indicating that country-level geographic labelling was internally consistent where location metadata were present. The geographic distribution of records was strongly concentrated among a small number of countries. The United States contributed 44,112 records (44.1%), followed by the Netherlands (17,690; 17.7%), France (11,588; 11.6%), and the United Kingdom (11,343; 11.3%). Together, these four countries accounted for 84.7% of all records, indicating substantial geographic concentration within the dataset Fig 4. Records labelled not provided (n = 381) had a mean vulnerability score of 12.0, the highest among all geographic label categories, reflecting the accumulation of missing metadata across multiple fields. Country-level vulnerability score distributions are summarised in Table 5.

**Figure 4:**
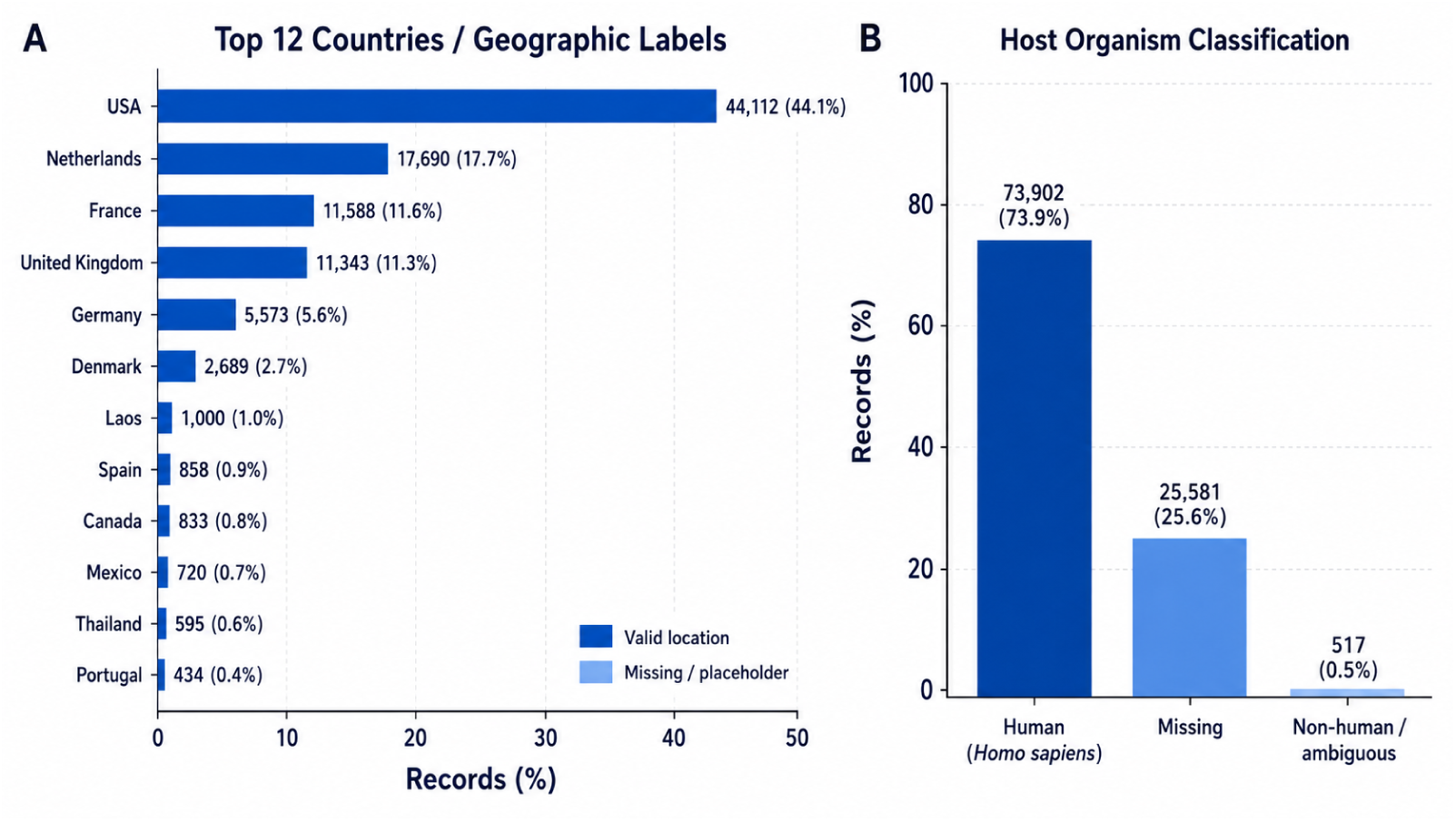
Geographic and host metadata analysis across 100,000 SARS-CoV-2 BioSample records. Left: top 12 country distribution, with the USA, Netherlands, France, and United Kingdom accounting for 84.7% of all records. Right: host organism classification, with 73.9% human, 25.6% missing, and 0.5% non-human or ambiguous.

**Table 5:**
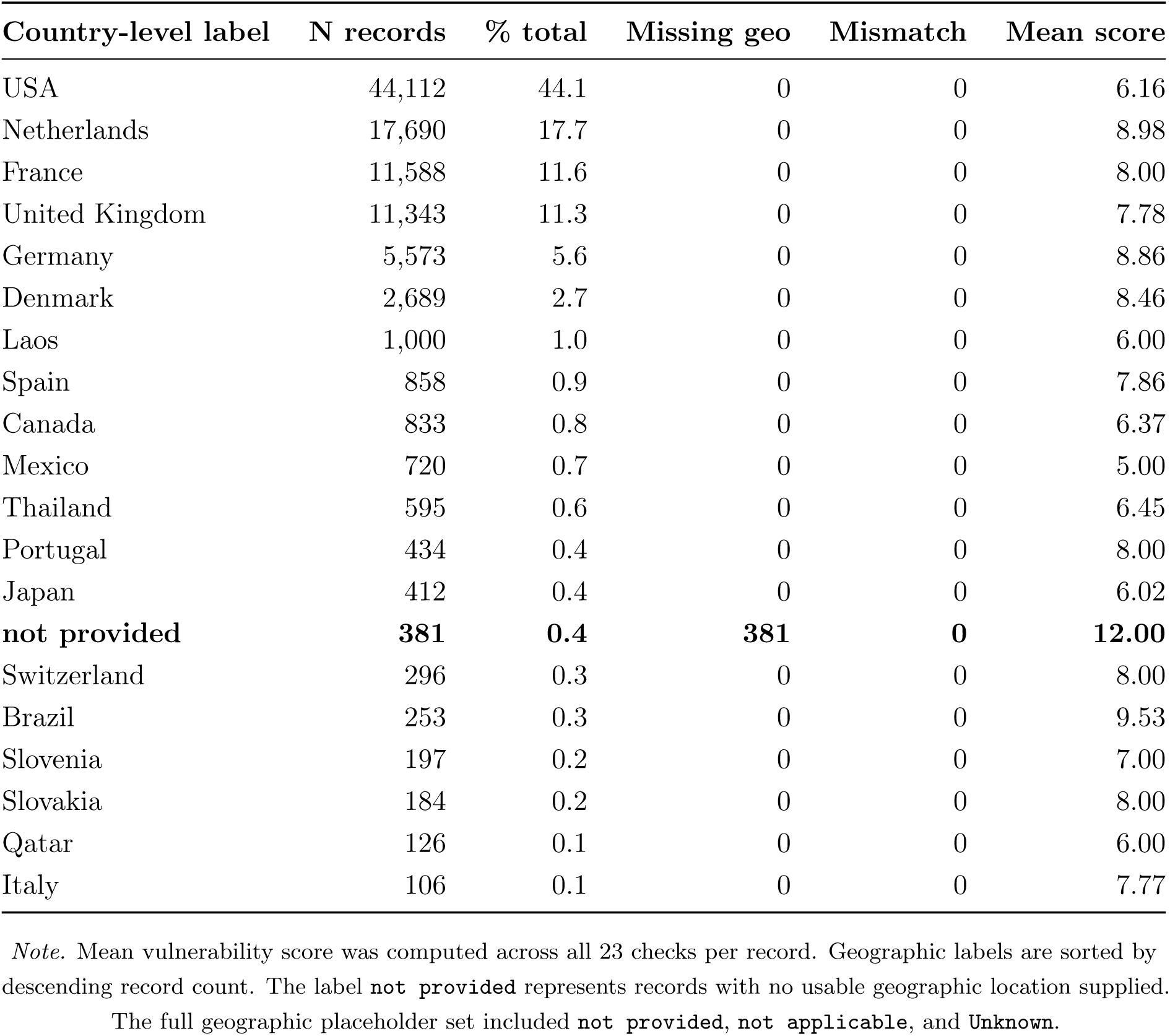
Country-level breakdown for the top 20 geographic labels, showing record counts, missing geography flags, country-field mismatches, and mean vulnerability scores. Records labelled not provided carried the highest mean vulnerability score (12.0).

| Country-level label | N records | % total | Missing geo | Mismatch | Mean score |
| --- | --- | --- | --- | --- | --- |
| USA | 44,112 | 44.1 | 0 | 0 | 6.16 |
| Netherlands | 17,690 | 17.7 | 0 | 0 | 8.98 |
| France | 11,588 | 11.6 | 0 | 0 | 8.00 |
| United Kingdom | 11,343 | 11.3 | 0 | 0 | 7.78 |
| Germany | 5,573 | 5.6 | 0 | 0 | 8.86 |
| Denmark | 2,689 | 2.7 | 0 | 0 | 8.46 |
| Laos | 1,000 | 1.0 | 0 | 0 | 6.00 |
| Spain | 858 | 0.9 | 0 | 0 | 7.86 |
| Canada | 833 | 0.8 | 0 | 0 | 6.37 |
| Mexico | 720 | 0.7 | 0 | 0 | 5.00 |
| Thailand | 595 | 0.6 | 0 | 0 | 6.45 |
| Portugal | 434 | 0.4 | 0 | 0 | 8.00 |
| Japan | 412 | 0.4 | 0 | 0 | 6.02 |
| **not provided** | **381** | **0.4** | **381** | **0** | **12.00** |
| Switzerland | 296 | 0.3 | 0 | 0 | 8.00 |
| Brazil | 253 | 0.3 | 0 | 0 | 9.53 |
| Slovenia | 197 | 0.2 | 0 | 0 | 7.00 |
| Slovakia | 184 | 0.2 | 0 | 0 | 8.00 |
| Qatar | 126 | 0.1 | 0 | 0 | 6.00 |
| Italy | 106 | 0.1 | 0 | 0 | 7.77 |

*Note.* Mean vulnerability score was computed across all 23 checks per record. Geographic labels are sorted by descending record count. The label **not provided** represents records with no usable geographic location supplied.
The full geographic placeholder set included **not provided**, **not applicable**, and **Unknown**.

##### Host and Specimen Metadata Layer

Host and specimen metadata vulnerability indicators were identified in 51,963 records (52.0%), making this the most affected biological metadata layer in the dataset. The most substantial contribution to this result was the absence of isolation source information: 50,908 records (50.9%) had missing or functionally missing isolation source metadata (H3; Table 6). The host organism field itself was missing or functionally missing in 25,581 records (25.6%) (H1; Table 6), limiting interpretation of transmission context and sample origin. Within the formal H2 audit check, 12 records were flagged by the conservative exact-match screen as non-human or laboratory-associated host contexts (Table 6), including records associated with VERO cell culture lines. These entries represent laboratory-propagated virus rather than primary host attribution. An additional 326 records (0.3%) contained biologically implausible isolation source entries (H5; Table 6), including values such as ‘library’ and ‘cultured virus’, which are more consistent with laboratory processing contexts than primary clinical sampling. The broader host classification shown in Figure 4 groups 517 records (0.5%) as non-human or ambiguous, since it includes free-text and inconsistently formatted host values. The formal H2 check applies a stricter exact-string screen, which is why it flags only the 12 records reported in Table 6. The difference reflects the conservatism of the audit check rather than a discrepancy in the underlying data.

**Table 6:** Individual audit check results across all 23 checks. Zero-flag checks indicate that extreme temporal, geographic, host, and specimen anomaly types were not observed in this dataset; dominant vulnerabilities were structural metadata absences rather than logical contradictions.

| ID | Layer | Description | N flagged | % flagged | N clean | % clean |
| --- | --- | --- | --- | --- | --- | --- |
| T1 | Temporal | Missing or unparseable collection date | 1,116 | 1.1 | 98,884 | 98.9 |
| T2 | Temporal | Year-only collection-date precision | 1,279 | 1.3 | 98,721 | 98.7 |
| T3 | Temporal | Month-year collection-date precision | 1,409 | 1.4 | 98,591 | 98.6 |
| T4 | Temporal | Pre-emergence collection date (<1 Nov 2019) | 0 | 0.0 | 100,000 | 100.0 |
| T5 | Temporal | Future collection date | 0 | 0.0 | 100,000 | 100.0 |
| T6 | Temporal | Collection date after submission date | 0 | 0.0 | 100,000 | 100.0 |
| T7 | Temporal | Same-day collection and submission | 0 | 0.0 | 100,000 | 100.0 |
| G1 | Geographic | Missing or functionally missing geographic location | 479 | 0.5 | 99,521 | 99.5 |
| G2 | Geographic | Country field inconsistent with `geographic_location` | 0 | 0.0 | 100,000 | 100.0 |
| G3 | Geographic | Geographic field contains placeholder text | 479 | 0.5 | 99,521 | 99.5 |
| H1 | Host | Missing or functionally missing host organism field | 25,581 | 25.6 | 74,419 | 74.4 |
| H2 | Host | Non-human or laboratory-associated host organism by exact-match screen | 12 | < 0.1 | 99,988 | > 99.9 |
| H3 | Host | Missing or functionally missing isolation source | 50,908 | 50.9 | 49,092 | 49.1 |
| H4 | Host | Isolation source placeholder text not already captured under H3 | 0 | 0.0 | 100,000 | 100.0 |
| H5 | Host | Implausible isolation source for a human respiratory virus | 326 | 0.3 | 99,674 | 99.7 |
| P1 | Provenance | Missing `collected_by` field | 51,348 | 51.3 | 48,652 | 48.7 |
| P2 | Provenance | Missing BioProject linkage within BioSample metadata | 99,280 | 99.3 | 720 | 0.7 |
| P3 | Provenance | Missing GenBank accession within BioSample metadata | 100,000 | 100.0 | 0 | 0.0 |
| P4 | Provenance | Missing SRA accession within BioSample metadata | 3,744 | 3.7 | 96,256 | 96.3 |
| A1 | Technical | Missing sequencing platform | 99,909 | 99.9 | 91 | 0.1 |
| A2 | Technical | Missing Pango lineage assignment | 98,139 | 98.1 | 1,861 | 1.9 |
| A3 | Technical | Missing coverage or sequencing depth | 100,000 | 100.0 | 0 | 0.0 |
| A4 | Technical | Missing assembly method | 100,000 | 100.0 | 0 | 0.0 |

*Note.* Percentages were calculated using the full dataset denominator of 100,000 records.

Together, these findings indicate substantial incompleteness in host and specimen metadata within the BioSample attribute layer. Such gaps are directly relevant to host and specimen metadata poisoning scenarios described in the APBT taxonomy, in which incorrect host attribution could distort interpretation of zoonotic transmission pathways or obscure the clinical origin of sequences within genomic surveillance datasets [46, 47, 61].

##### Provenance and Administrative Metadata Layer

The completeness of provenance linkage fields and analytical metadata attributes within the BioSample layer is summarised in Fig. 5. Provenance and administrative metadata vulnerability indicators were identified in all 100,000 records (100.0%), primarily because at least one provenance field was absent in every record (Table 6). GenBank accession numbers (P3) were absent from the BioSample metadata layer for all records in the dataset (100,000 records; 100.0%; Table 6). This reflects a structural feature of the BioSample submission model in which nucleotide sequence accessions are typically assigned through a separate submission workflow and are not consistently cross-referenced at the BioSample level [60, 68]. BioProject linkage (P2) was absent in 99,280 records (99.3%), with only 720 records (0.7%) carrying a BioProject accession within the BioSample metadata layer (Table 6). The collected_by field (P1), which provides institutional attribution for sample collection, was missing in 51,348 records (51.3%). SRA accession linkage (P4) was absent in 3,744 records (3.7%), while 96,256 records (96.3%) had an SRA accession visible within the BioSample metadata layer (Table 6).

**Figure 5:**
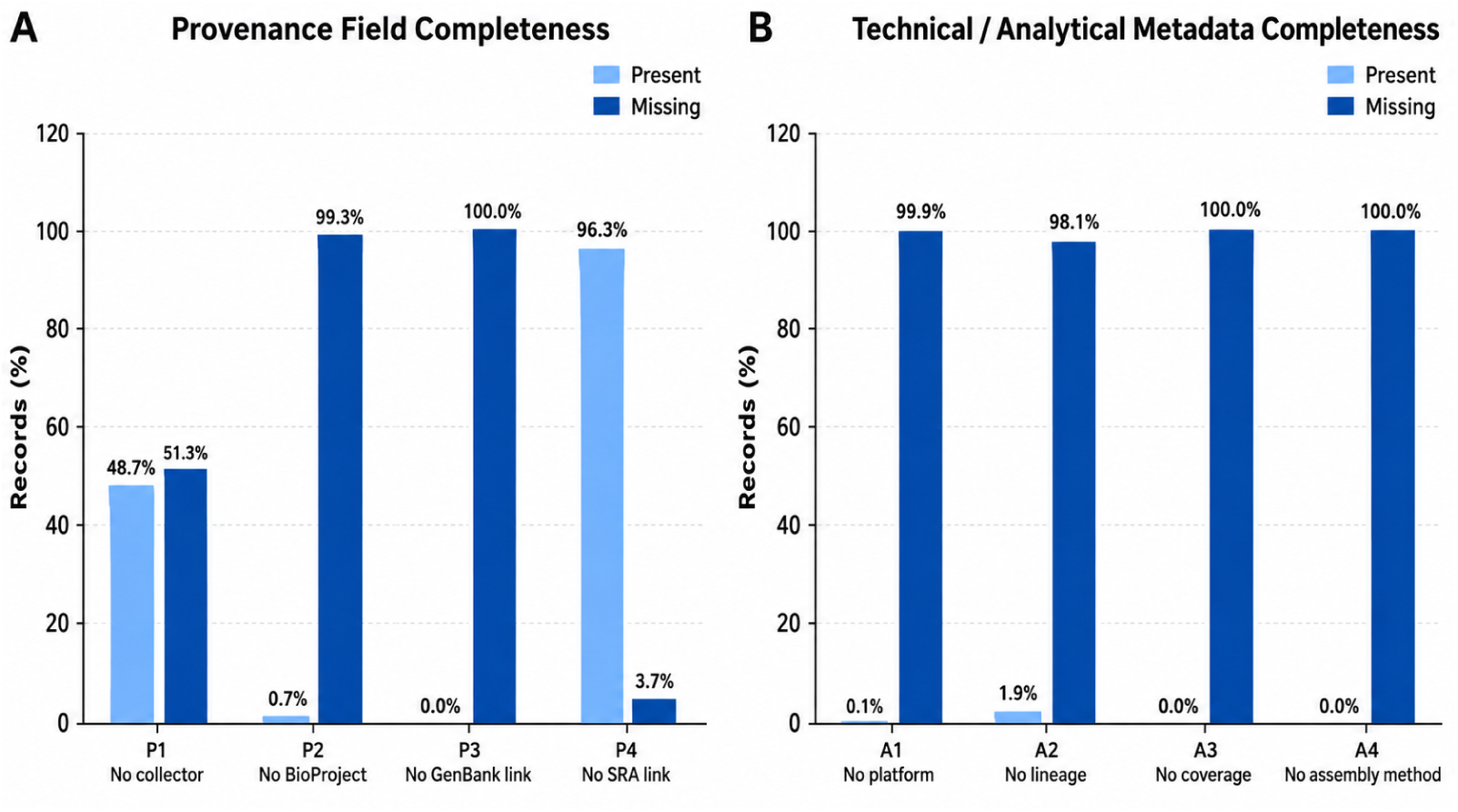
Provenance and technical metadata completeness across 100,000 SARS-CoV-2 BioSample records. Left: provenance fields P1–P4, with GenBank linkage (P3) absent from 100.0% of records and BioProject linkage (P2) from 99.3%. Right: technical and analytical fields A1–A4, showing near-universal absence of sequencing platform, coverage depth, assembly method, and Pango lineage.

These provenance findings are particularly relevant in the context of APBT Phase 2 infiltration detection. The absence of BioProject linkage in 99.3% of records limits the ability to contextualise individual BioSample submissions within broader project-level submission frameworks, reducing the visibility of anomalous submission patterns at the repository metadata level [48]. Similarly, the absence of GenBank cross-references within BioSample metadata means that the nucleotide sequence associated with each BioSample record cannot be directly verified from the BioSample record alone. Together, these gaps indicate substantial limitations in chain-of-custody reconstruction using BioSample metadata attributes in isolation [64, 69].

##### Technical and Analytical Metadata Layer

Technical and analytical metadata vulnerability indicators were identified in all 100,000 records (100.0%), making this, alongside provenance metadata, one of the two universally affected metadata layers in the dataset (Table 6; Fig. 5). Coverage depth (A3) and assembly method (A4) were absent from all 100,000 records. Sequencing platform information (A1) was absent from 99,909 records (99.9%), with only 91 records (0.1%) providing platform metadata. Pango lineage classification (A2), a widely used analytical designation in SARS-CoV-2 variant surveillance workflows, was absent from 98,139 records (98.1%), with only 1,861 records (1.9%) containing lineage assignments (Table 6) [50, 55]. The near-universal absence of these technical attributes reflects a structural feature of the BioSample metadata model, in which sequencing platform information, analytical lineage classification, coverage depth, and assembly method are not mandatory components of the BioSample attribute schema and may instead be recorded in linked repositories or downstream analytical resources such as GenBank, SRA, or lineage-assignment workflows [60, 68, 70]. As a result, analytical provenance cannot be independently verified using BioSample metadata attributes alone.

These findings are directly relevant to the analytical metadata exposure surface described in the APBT taxonomy, in which incomplete analytical context increases the difficulty of detecting inconsistencies between sequence interpretation, lineage designation, and sequencing methodology at the repository metadata level [62, 65, 66].

##### Submitter Concentration Analysis

Analysis of submitter concentration revealed substantial dominance of the dataset by a small number of contributing institutions, a pattern directly relevant to dataset concentration exposure surfaces within the APBT framework. The institutional concentration profile of dataset submissions is illustrated in Fig 6. A single institution, the European Bioinformatics Institute (EBI), contributed 50,231 records (50.2%) of the entire dataset and exhibited a mean vulnerability score of 8.50, the highest among major submitting institutions [58, 60]. Concentration curve analysis further showed that a single submitting laboratory accounted for approximately half of all records in the dataset, while the ten largest submitting institutions together accounted for approximately 80% of all records. This level of institutional concentration indicates that the aggregate vulnerability profile of the dataset is strongly influenced by a relatively small number of contributors. Such concentration has important implications for repository-level robustness. Systematic metadata irregularities affecting a major submitting institution could influence a substantial proportion of records available for downstream genomic surveillance analyses [71, 72].

**Figure 6:**
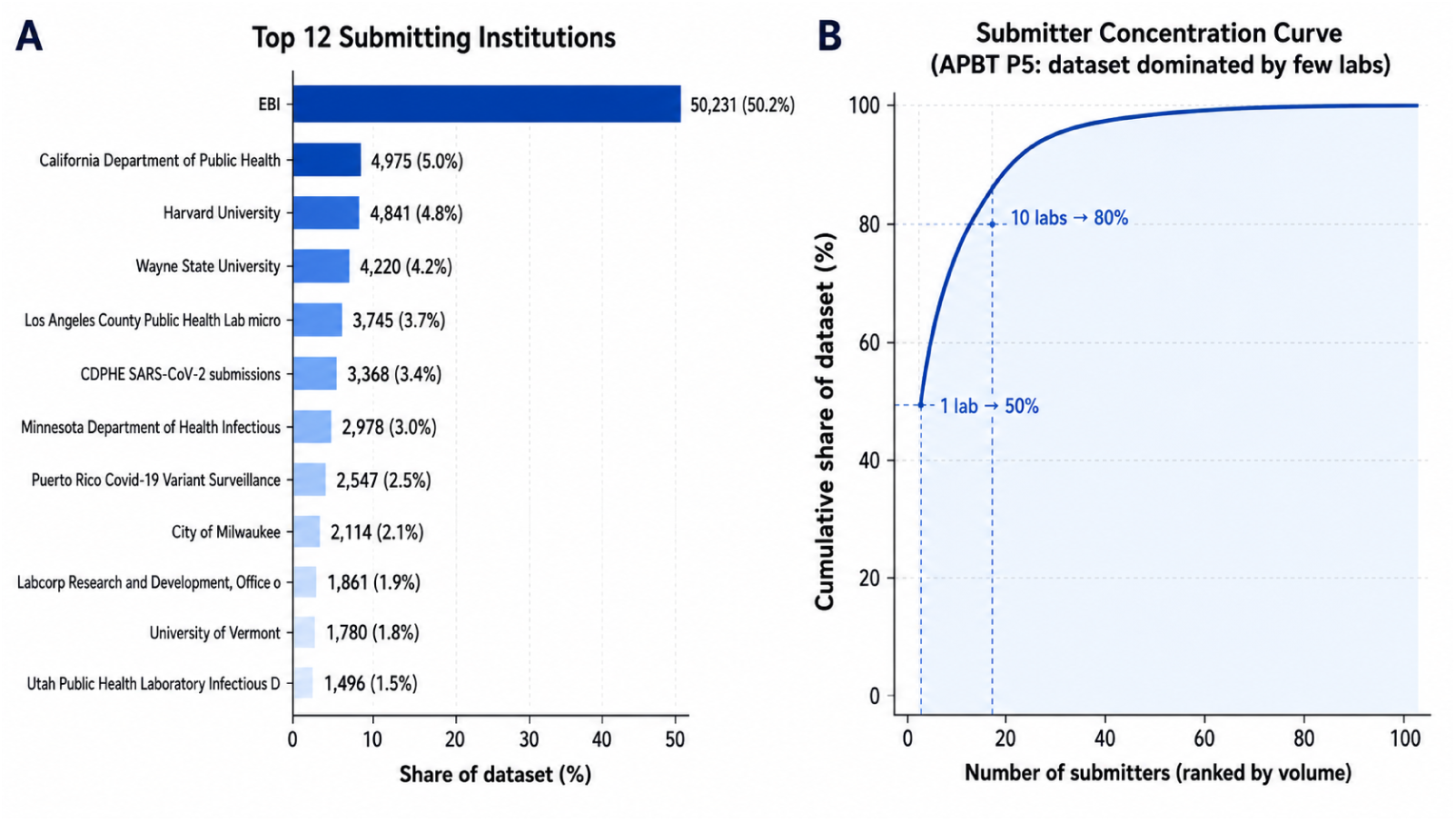
Submitter concentration analysis across 100,000 SARS-CoV-2 BioSample records. Left: top 12 submitting institutions by share of the dataset, with EBI alone accounting for 50.2%. Right: cumulative concentration curve, showing that the 10 largest institutions together account for approximately 80% of all records.

These findings highlight the importance of considering submission-level concentration patterns when evaluating metadata integrity exposure surfaces within large-scale genomic repositories. Table 7 presents the full submitter-level analysis for the top 12 contributing institutions.

**Table 7:**
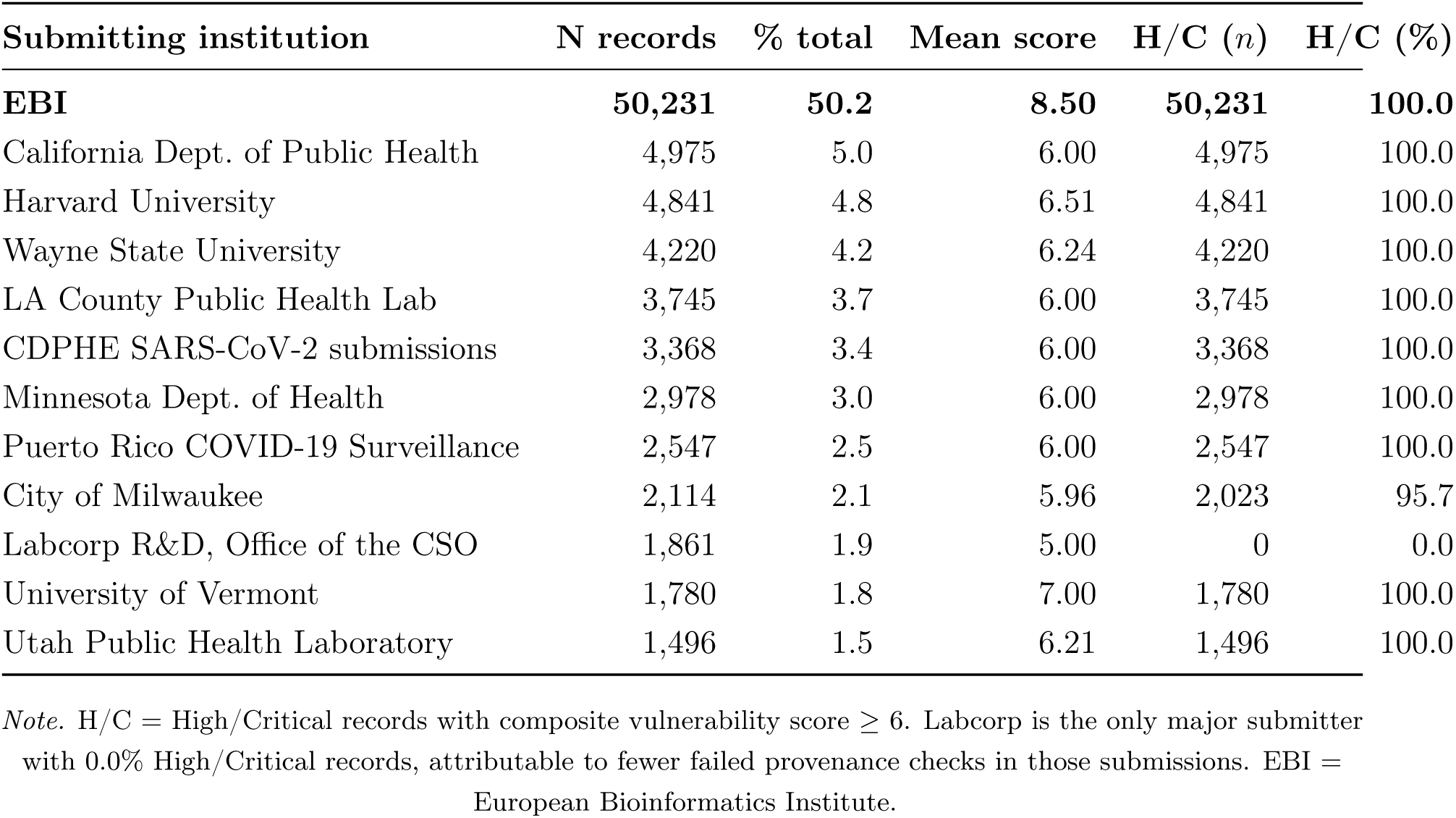
Top 12 submitting institutions ranked by record volume. Mean vulnerability score and proportion of High/Critical records are shown for each institution. EBI contributes 50.2% of records, with 100.0% classified in the High/Critical category.

### 3.3 Discussion

Our findings indicate that APBT-relevant metadata vulnerabilities are not isolated data-quality problems, but systematic features of the SARS-CoV-2 BioSample metadata layer. All 100,000 audited records carried at least one APBT-relevant metadata vulnerability indicator. On average, records triggered 7.34 out of 23 possible checks (SD 1.34), and 97.3% were classified as High or Critical severity. Exposure on this scale cannot be attributed to occasional submission errors, and it is consistent with earlier evidence that repository metadata are systematically uneven; Gonçalves et al. [7], auditing millions of BioSample records, reported that attribute names and values are largely uncontrolled and inconsistently represented. What we add to that picture is a shift of interpretation, since we read the same incompleteness not only as a barrier to reuse but also as an adversarial exposure surface.

The exposure was not distributed uniformly across the five metadata layers, and this distribution proves more informative than the aggregate rate. Temporal and geographic metadata were largely complete, with indicators in only 3.8% and 0.5% of records respectively, which indicates that submitters generally supply a usable collection date and a country-level location label. The provenance and technical layers showed the reverse pattern, since every record was affected. GenBank accession linkage was absent throughout, and BioProject linkage was missing from 99.3% of records. The technical fields were almost as sparse: sequencing platform, coverage depth, assembly method and Pango lineage were each absent from between 98.1% and 100.0% of records. In short, the fields that describe a sample are comparatively well kept, whereas the fields that would allow it to be verified are largely absent. For an adversarial model this is the more concerning ordering, since provenance and technical fields are precisely the metadata elements that could help expose a poisoned, synthetic or misattributed record.

Notably, none of the logical integrity checks were triggered. We found no pre-emergence collection dates, no future-dated collections, no records collected after their own submission, and no mismatches between the geographic and country fields. We therefore find no evidence of obvious or logically detectable manipulation in this snapshot, and the vulnerability we document is primarily one of incompleteness rather than demonstrated falsification. This distinction matters for how the results should be read, because an obvious contradiction can be detected and removed, whereas a well-formed record that simply lacks provenance is indistinguishable from the many legitimate records around it that also lack it. The weakness we identify therefore sits one layer earlier than the pipeline and workflow vulnerabilities reported in prior work [16, 17], at the point where a record enters the repository rather than where it is subsequently processed.

These gaps have concrete consequences for the scenarios the APBT framework anticipates. Where an adversary seeks to introduce a misleading record, detection depends on institutional and project-level context, yet BioProject linkage is missing from 99.3% of records and collected_by from 51.3%. When most legitimate records already lack traceable provenance, its absence no longer functions as a warning sign, and a fabricated record carrying a plausible organism, date and country would not stand out. The submitter distribution reinforces this concern, since half the dataset (50.2%) enters through a single handle, EBI, and the ten largest contributors account for roughly 80%. EBI is largely a brokering route for records mirrored from ENA rather than a single laboratory, but the practical point holds: the conventions of a few high-volume channels shape the completeness of the whole repository. The near-total absence of technical fields extends the same problem to records that persist over time, because without platform, coverage or assembly information a record cannot be cross-checked for technical plausibility using BioSample metadata alone. As large-scale surveillance relies on aggregation rather than manual inspection, systematic gaps of this kind can quietly weaken automated quality control across many records at once.

Taken together, these observations point to a practical tension for open repositories rather than a case for restriction. Repositories such as NCBI BioSample are deliberately permissive so that data can circulate quickly during an emergency, and that openness is worth preserving. A proportionate response is tiered metadata assurance, in which basic submission remains open while records intended for forensic or policy-relevant use are asked to carry stronger provenance and technical metadata, with priority on BioProject linkage, collected_by, sequencing platform, coverage depth, assembly method and lineage. Because a small number of submitters dominate the dataset, targeted feedback and completeness reporting for those contributors would raise repository-wide integrity at modest cost, without burdening the long tail of low-volume laboratories.

Several limitations qualify these conclusions. We audited metadata as exposed through the BioSample attribute model, so the complete absence of fields such as GenBank accession and assembly method reflects the scope of the BioSample record rather than the total absence of that information across linked resources such as GenBank, SRA or ENA [58]; the 100% figures should therefore be read as completeness limitations of the BioSample metadata layer, not of the sequence records in their entirety. The checks are binary and the composite score is additive and unweighted, so a record missing several minor fields can score much like one missing a few critical fields, and the severity tiers are best interpreted as cumulative exposure rather than as a validated risk metric. Most importantly for the framework, the audit detects missing values, placeholders and logical contradictions, but not manipulation that is internally consistent, since a fabricated record that is plausible in every field could pass every check. That is the scenario the APBT framework treats as most dangerous, and the one metadata auditing alone cannot resolve. Finally, our analysis covers a single snapshot. The audited records form a registration ordered window rather than a random sample, and this shapes part of the profile we report. A window of this kind captures complete submission batches from the institutions most active during the corresponding period, so the submitter concentration described in Section 3.2.2 should be read as a property of this window rather than as a stable characteristic of BioSample as a whole. The layer level completeness findings are less sensitive to the sampling design, since the fields most often absent are optional under the BioSample schema for every submitter, but a randomised or year stratified draw would be required to confirm that the observed rates hold across the full repository.

## 4 Conclusion & Future directions

This paper introduced APBTs as a framework for understanding how malicious actors could degrade genomic surveillance through data centric manipulation. We presented a multilayer APBT taxonomy organised around a four phase adversarial lifecycle and operationalised this framework through an empirical audit of 100,000 SARS-CoV-2 BioSample records from NCBI. The findings demonstrate that APBT relevant metadata vulnerabilities are systematically present at repository scale. Every record triggered at least one vulnerability indicator, with a mean of 7.34 per record, and 97.3 percent fell within High or Critical severity categories. While temporal and geographic metadata were comparatively complete, provenance and technical metadata were universally affected. GenBank linkage was absent from all records, BioProject linkage from 99.3 percent, and technical fields including sequencing platform, coverage depth, and assembly method were almost entirely missing. This asymmetry creates conditions where fabricated records could evade detection. No logical inconsistencies were detected, confirming that vulnerabilities are structural incompleteness rather than demonstrated falsification.

Several limitations point toward future research. Examining whether incompleteness persists across linked repositories including GenBank, SRA, and ENA would establish whether cross repository linkage enables provenance reconstruction. Developing methods for detecting plausible but fabricated records beyond format validation is essential, as consistent fabrications would pass all current checks. Comparative audits across pathogens, repositories, and time periods would establish generalisability. Validating vulnerability scores through red team exercises and controlled poisoning experiments would identify which metadata gaps are most consequential. Tracing provenance through full submission chains would reveal where provenance is most vulnerable.

Future work should also examine how APBTs might interact with AI enabled sequence generation and synthetic biology, and develop quantitative risk assessment frameworks. Governance questions including balancing openness against security, international coordination mechanisms, and maintaining trust when integrity gaps are identified also merit investigation. The COVID-19 pandemic demonstrated both the power of open genomic sharing and the vulnerability of systems that depend on trust alone. As genomic surveillance expands and AI accelerates plausible sequence generation, verifiable data infrastructure becomes essential. The question is whether the community will address these vulnerabilities before they are exploited.

## 5 Other Information

## Acknowledgements

Authors acknowledge the use of generative AI tools to assist with language editing, to improve the clarity and structure of the manuscript text, to enhance the presentation quality of the figures for publication, and to help debug the code. All research design, data, analysis, results, and interpretation are the authors’ own. The authors reviewed and verified all AI-assisted content, and take full responsibility for the content of the publication.

## AUTHOR CONTRIBUTIONS

NA: Conceptualization, Methodology, Software, Formal analysis, Investigation, Data curation, Visualization, Original draft preparation, Review and editing, Supervision, Project administration. MK: Methodology, Investigation, Original draft preparation, Review and editing. HAA: Validation, Investigation, Review and editing, Funding acquisition. AK: Methodology, Validation, Review and editing. MuH: Validation, Investigation, Review and editing. BJW: Validation, Review and editing. All authors contributed to the article and approved the submitted version.

## DATA AVAILABILITY STATEMENT

The data analysed in this study are publicly available from the NCBI BioSample repository (https://www.ncbi.nlm.nih.gov/biosample). Records were retrieved in read only mode using the NCBI Entrez API with the taxonomy identifier txid2697049[Organism]. No repository records were modified. The derived audit dataset, the 23 check audit framework, and the analysis code used to generate all results and figures are available from the corresponding author on reasonable request.

## ETHICS STATEMENT

Ethical approval was not required for this study. The research used only publicly available metadata records held in the NCBI BioSample repository. No human participants, animals, or biological samples were involved, and no personally identifiable information was accessed or analysed. All data were retrieved in read only mode through publicly documented interfaces in accordance with the repository’s published usage guidelines.

## Footnotes

1 A genomic sequence is a digital representation of DNA or RNA, typically stored in formats such as FASTA, FASTQ, SAM, or BAM. These files contain the nucleotide content of a sample and serve as the foundation for downstream analyses including alignment, variant detection, phylogenetic reconstruction, and ML [44].

## References

[1] Ben Hu, Hua Guo, Peng Zhou, and Zheng-Li Shi. Characteristics of sars-cov-2 and covid-19. Nature reviews microbiology, 19(3):141–154, 2021. doi: 10.1038/s41579-020-00459-7.

[2] Nasreen Anjum, Hani Alshahrani, Darakhshan Syed, Asadullah Shaikh, and Mahreen Ul Hassan. Next generation sequencing under attack: investigating insider threats and organizational behaviour. PeerJ Computer Science, 11:e3008, 2025. doi: 10.7717/peerj-cs.3008.

[3] Talha Burki. First shared sars-cov-2 genome: Gisaid vs virological. org. The Lancet Microbe, 4(6):e395, 2023.

[4] Anderson F Brito, Elizaveta Semenova, Gytis Dudas, Gabriel W Hassler, Chaney C Kalinich, Moritz UG Kraemer, Joses Ho, Houriiyah Tegally, George Githinji, Charles N Agoti, et al. Global disparities in sars-cov-2 genomic surveillance. Nature communications, 13(1):7003, 2022. doi: 10.1038/s41467-022-33713-y.

[5] Shruti Khare, Céline Gurry, Lucas Freitas, Mark B Schultz, Gunter Bach, Amadou Diallo, Nancy Akite, Joses Ho, Raphael TC Lee, Winston Yeo, et al. Gisaid’s role in pandemic response. China CDC weekly, 3(49):1049, 2021.

[6] Matteo Chiara, Anna Maria D’Erchia, Carmela Gissi, Caterina Manzari, Antonio Parisi, Nicoletta Resta, Federico Zambelli, Ernesto Picardi, Giulio Pavesi, David S. Horner, et al. Next generation sequencing of SARS-CoV-2 genomes: challenges, applications and opportunities. Briefings in Bioinformatics, 22(2):616–630, 2021. doi: 10.1093/bib/bbaa297.

[7] Rafael S Gonçalves and Mark A Musen. The variable quality of metadata about biological samples used in biomedical experiments. Scientific data, 6(1):190021, 2019.

[8] James Diggans and Emily Leproust. Next steps for access to safe, secure dna synthesis. Frontiers in bioengineering and biotechnology, 7:86, 2019. doi: 10.3389/fbioe.2019.00086.

[9] David A. Alber, Zane Yang, Anton Alyakin, Eric Yang, Sujay Rai, Aly A. Valliani, et al. Medical large language models are vulnerable to data-poisoning attacks. Nature Medicine, 31(2):618–626, 2025. doi: 10.1038/s41591-024-03445-1.

[10] Ibrahim Ghafir, Vaclav Prenosil, et al. Advanced persistent threat attack detection: an overview. Int J Adv Comput Netw Secur, 4(4):5054, 2014.

[11] Sandro Quintero-Bonilla and Angel Martín del Rey. A new proposal on the advanced persistent threat: a survey. Applied Sciences, 10(11):3874, 2020. doi: 10.3390/app10113874.

[12] Noran Shafik Fouad. Cyberbiosecurity in the new normal: Cyberbio risks, pre-emptive security, and the global governance of bioinformation. European Journal of International Security, 9(4):553–573, 2024. doi: 10.1017/eis.2024.19.

[13] Eleonore Pauwels. Cyber-biosecurity: How to protect biotechnology from adversarial AI attacks. Hybrid CoE Strategic Analysis 26, European Centre of Excellence for Countering Hybrid Threats (Hybrid CoE), Helsinki, Finland, May 2021. URL https://www.hybridcoe.fi/wp-content/uploads/2021/05/20210503_Hybrid_CoE_Strategic_Analysis_26_Cyber_biosecurity_WEB.pdf.

[14] Nasreen Anjum, Hani Alshahrani, Asadullah Shaikh, Mahreen Kiran, Shah Raz, Abu Alam, et al. Cyber-biosecurity challenges in next-generation sequencing: A comprehensive analysis of emerging threat vectors. IEEE Access, 13:52006–52035, 2025. doi: 10.1109/ACCESS.2025.3552069.

[15] Boris A Vinatzer, Lenwood S Heath, Hussain MJ Almohri, Michael J Stulberg, Christopher Lowe, and Song Li. Cyberbiosecurity challenges of pathogen genome databases. Frontiers in bioengineering and biotechnology, 7:106, 2019. doi: 10.3389/fbioe.2019.00106.

[16] Garrett J Schumacher, Sterling Sawaya, Demetrius Nelson, and Aaron J Hansen. Genetic information insecurity as state of the art. Frontiers in bioengineering and biotechnology, 8: 591980, 2020. doi: 10.3389/fbioe.2020.591980.

[17] Jacob Caswell, Jason D Gans, Nicholas Generous, Corey M Hudson, Eric Merkley, Curtis Johnson, Christopher Oehmen, Kristin Omberg, Emilie Purvine, Karen Taylor, et al. Defending our public biological databases as a global critical infrastructure. Frontiers in bioengineering and biotechnology, 7:58, 2019. doi: 10.3389/fbioe.2019.00058.

[18] Peter Ney, Karl Koscher, Lee Organick, Luis Ceze, and Tadayoshi Kohno. Computer security, privacy, and {DNA} sequencing: compromising computers with synthesized {DNA}, privacy leaks, and more. In 26th USENIX Security Symposium (USENIX Security 17), pages 765–779, 2017.

[19] Peter Martin Ney. Securing the Future of Biotechnology: A Study of Emerging Bio-Cyber Security Threats to DNA-Information Systems. PhD thesis, University of Washington, 2019. URL http://hdl.handle.net/1773/44143.

[20] Ana León, José Reyes, Verónica Burriel, and Francisco Valverde. Data quality problems when integrating genomic information. In International Conference on Conceptual Modeling, pages 173–182. Springer, 2016.

[21] Farah Zaib Khan, Stian Soiland-Reyes, Richard O Sinnott, Andrew Lonie, Carole Goble, and Michael R Crusoe. Sharing interoperable workflow provenance: A review of best practices and their practical application in cwlprov. GigaScience, (11):giz095, 2019. doi: 10.1093/gigascience/giz095.

[22] Suyash S Shringarpure and Carlos D Bustamante. Privacy risks from genomic data-sharing beacons. The American Journal of Human Genetics, 97(5):631–646, 2015. doi: 10.1016/j.ajhg.2015.09.010.

[23] Zaixi Zhang, Zhenghong Zhou, Ruofan Jin, Le Cong, and Mengdi Wang. Genebreaker: Jailbreak attacks against dna language models with pathogenicity guidance. arXiv preprint arXiv:2505.23839, 2025.

[24] Nathan Killoran, Leo J Lee, Andrew Delong, David Duvenaud, and Brendan J Frey. Generating and designing dna with deep generative models. arXiv preprint arXiv:1712.06148, 2017.

[25] Roger Brent and T Greg McKelvey Jr. Contemporary ai foundation models increase biological weapons risk. *arXiv preprint arXiv:2506.13798*, 2025.

[26] National Institute of Standards and Technology. Biosecurity for synthetic nucleic acid sequences. Technical report, National Institute of Standards and Technology, 2024. URL https://www.nist.gov/programs-projects/biosecurity-synthetic-nucleic-acid-sequences. Accessed: 2026.

[27] UK AI Safety Institute. Frontier ai safety and biosecurity evaluation. Technical report, UK Department for Science Innovation and Technology, 2024.

[28] Saurabh Ranjan, Lucas Potter, and Xavier-Lewis Palmer. A snapshot of the biocybersecurity/cyberbiosecurity landscape. In Proceedings of the 19th International Conference on Cyber Warfare and Security. Academic Conferences and publishing limited, 2025.

[29] Patrick F Walsh. Securing the bioeconomy: Exploring the role of cyberbiosecurity. In The handbook of security, pages 335–355. Springer, 2022.

[30] Richard Annan, Justin Noland, Kamaria Perkins, Xiaohong Yuan, Kaushik Roy, and Letu Qingge. Genomic privacy and security in the era of artificial intelligence and quantum computing. Discover Computing, 28(1):108, 2025. doi: 10.1007/s10791-025-09627-w.

[31] Samuel G Finlayson, John D Bowers, Joichi Ito, Jonathan L Zittrain, Andrew L Beam, and Isaac S Kohane. Adversarial attacks on medical machine learning. Science, 363(6433): 1287–1289, 2019.

[32] Avi Schwarzschild, Micah Goldblum, Arjun Gupta, John P Dickerson, and Tom Goldstein. Just how toxic is data poisoning? a unified benchmark for backdoor and data poisoning attacks. In Proceedings of the 38th International Conference on Machine Learning, volume 139 of Proceedings of Machine Learning Research, pages 9389–9398. PMLR, 2021.

[33] Aniruddha Saha, Akshayvarun Subramanya, and Hamed Pirsiavash. Hidden trigger backdoor attacks. In Proceedings of the AAAI conference on artificial intelligence, volume 34, pages 11957–11965, 2020. doi: 10.1609/aaai.v34i07.6871.

[34] Yuezun Li, Yiming Li, Baoyuan Wu, Longkang Li, Ran He, and Siwei Lyu. Invisible backdoor attack with sample-specific triggers. In Proceedings of the IEEE/CVF international conference on computer vision, pages 16463–16472, 2021.

[35] Jonas Geiping, Hartmut Bauermeister, Hannah Dröge, and Michael Moeller. Inverting gradients-how easy is it to break privacy in federated learning? Advances in neural information processing systems, 33:16937–16947, 2020.

[36] Hao Fang, Yixiang Qiu, Hongyao Yu, Wenbo Yu, Jiawei Kong, Baoli Chong, Bin Chen, Xuan Wang, Shu-Tao Xia, and Ke Xu. Privacy leakage on dnns: A survey of model inversion attacks and defenses. *arXiv preprint arXiv:2402.04013*, 2024.

[37] Wencheng Yang, Song Wang, Di Wu, Taotao Cai, Yanming Zhu, Shicheng Wei, Yiying Zhang, Xu Yang, Zhaohui Tang, and Yan Li. Deep learning model inversion attacks and defenses: a comprehensive survey. Artificial Intelligence Review, 58(8):242, 2025. doi: 10.1007/s10462-025-11248-0.

[38] Jie Lu, Anjin Liu, Fan Dong, Feng Gu, Joao Gama, and Guangquan Zhang. Learning under concept drift: A review. IEEE transactions on knowledge and data engineering, 31(12): 2346–2363, 2018. doi: 10.1109/TKDE.2018.2876857.

[39] Adriana Sayuri Iwashita and Joao Paulo Papa. An overview on concept drift learning. IEEE access, 7:1532–1547, 2018. doi: 10.1109/ACCESS.2018.2886026.

[40] Charalampos Koilakos, Ioannis Mouratidis, and Ilias Georgakopoulos-Soares. Poisoning the genome: Targeted backdoor attacks on dna foundation models. arXiv preprint arXiv:2603.27465, 2026.

[41] Pelin Yilmaz, Renzo Kottmann, Dawn Field, Rob Knight, James R Cole, Linda Amaral-Zettler, Jack A Gilbert, Ilene Karsch-Mizrachi, Anjanette Johnston, Guy Cochrane, et al. Minimum information about a marker gene sequence (mimarks) and minimum information about any (x) sequence (mixs) specifications. Nature biotechnology, 29(5):415–420, 2011.

[42] Emma J Griffiths, Ruth E Timme, Catarina Inês Mendes, Andrew J Page, Nabil-Fareed Alikhan, Dan Fornika, Finlay Maguire, Josefina Campos, Daniel Park, Idowu B Olawoye, et al. Future-proofing and maximizing the utility of metadata: The PHA4GE SARS-CoV-2 contextual data specification package. GigaScience, 11:giac003, 2022. doi: 10.1093/gigascience/giac003.

[43] Shangzhe Li, Yue Shi, Jing Yang, Haizhou Liu, Lijia Jia, and Di Liu. Exploring potential biosafety implications in dna information storage. Biosafety and Health, 7(02):132–139, 2025.

[44] AJ Marian. Sequencing your genome: what does it mean? Methodist DeBakey cardiovascular journal, 10(1):3, 2014.

[45] European Genome-phenome Archive. What is metadata? EGA Documentation, 2025. URL https://ega-archive.org/submission/metadata/what-is-metadata/. Accessed: 2026.

[46] Oana Stroe. Improving metadata for genomic sequences. ENA Documentation, 2024. URL https://ena-docs.readthedocs.io/en/latest/faq/pathogen-subs-guide.html. Accessed: 2026.

[47] Kun Zhao, Katie Farrell, Melchizedek Mashiku, Dawit Abay, Kevin Tang, M Steven Oberste, and Cara C Burns. A search-based geographic metadata curation pipeline to refine sequencing institution information and support public health. Frontiers in Public Health, 11:1254976, 2023. doi: 10.3389/fpubh.2023.1254976.

[48] Bofeng Pan, Natalia Stakhanova, and Suprio Ray. Data provenance in security and privacy. ACM Computing Surveys, 55(14s):1–35, 2023. doi: 10.1145/3593294.

[49] NCBI Resource Coordinators. Homo sapiens genome assembly grch38 (gcf_000001405.26). https://www.ncbi.nlm.nih.gov/datasets/genome/GCF_000001405.26/, 2025. Accessed: 01-Dec-2025.

[50] cov-lineages.org. SARS-CoV-2 Pango lineage nomenclature and resources. https://cov-lineages.org/, 2025. Accessed: April 2026.

[51] Xin Yao, Tianchi Huang, Rui-Xiao Zhang, Ruiyu Li, and Lifeng Sun. Federated learning with unbiased gradient aggregation and controllable meta updating. *arXiv preprint arXiv:1910.08234*, 2019.

[52] Eugene Bagdasaryan, Andreas Veit, Yiqing Hua, Deborah Estrin, and Vitaly Shmatikov. How to backdoor federated learning. In Proceedings of the Twenty Third International Conference on Artificial Intelligence and Statistics, volume 108 of Proceedings of Machine Learning Research, pages 2938–2948, 2020.

[53] Vaishali Singh and SK Pandey. Revisiting cloud security attacks: Credential attack. In Rising Threats in Expert Applications and Solutions: Proceedings of FICR-TEAS 2020, pages 339–350. Springer, 2020.

[54] Abdul Razaque, Nazerke Shaldanbayeva, Bandar Alotaibi, Munif Alotaibi, Akhmetov Murat, and Aziz Alotaibi. Big data handling approach for unauthorized cloud computing access. Electronics, 11(1):137, 2022.

[55] James Hadfield, Colin Megill, Sidney M Bell, John Huddleston, Barney Potter, Charlton Callender, Pavel Sagulenko, Trevor Bedford, and Richard A Neher. Nextstrain: real-time tracking of pathogen evolution. Bioinformatics, 34(23):4121–4123, 2018. doi: 10.1093/bioinformatics/bty407.

[56] World Health Organization. Who covid-19 dashboard. https://data.who.int/dashboards/covid19/summary, 2025. Accessed: 03-Dec-2025.

[57] Ilene Karsch-Mizrachi, Toshihisa Takagi, Guy Cochrane, and International Nucleotide Sequence Database Collaboration. The international nucleotide sequence database collaboration. Nucleic Acids Research, 46(D1):D48–D51, 2018.

[58] Ilene Karsch-Mizrachi, Masanori Arita, Tony Burdett, Guy Cochrane, Yasukazu Nakamura, Kim D Pruitt, Valerie A Schneider, and on behalf of the International Nucleotide Sequence Database Collaboration. The international nucleotide sequence database collaboration (insdc): enhancing global participation. Nucleic acids research, 53(D1):D62–D66, 2025.

[59] National Center for Biotechnology Information. Entrez programming utilities help. https://www.ncbi.nlm.nih.gov/books/NBK25501/, 2024. Accessed: 2026.

[60] Dipayan Gupta, Isuru Liyanage, Yasset Perez-Riverol, Enrique Sapena Ventura, Wei Kheng Teh, Tim Van Den Bossche, Senthilnathan Vijayaraja, Amnon Khen, Ugis Sarkans, Tony Burdett, et al. Biosamples database: the global hub for sample metadata and multi-omics integration. Nucleic Acids Research, 54(D1):D1761–D1767, 2026. doi: 10.1093/nar/gkaf1133.

[61] Eric D Crandall, Rachel H Toczydlowski, Libby Liggins, Ann E Holmes, Maryam Ghoojaei, Michelle R Gaither, Briana E Wham, Andrea L Pritt, Cory Noble, Tanner J Anderson, et al. Importance of timely metadata curation to the global surveillance of genetic diversity. Conservation Biology, 37(4):e14061, 2023. doi: 10.1111/cobi.14061.

[62] Maximilian Sprang, Matteo Krüger, Miguel A Andrade-Navarro, and Jean-Fred Fontaine. Statistical guidelines for quality control of next-generation sequencing techniques. Life Science Alliance, 4(11), 2021. doi: 10.26508/lsa.202101113.

[63] World Health Organization. Global genomic surveillance strategy for pathogens with pandemic and epidemic potential 2022–2032. Technical report, World Health Organization, 2022. URL https://www.who.int/publications/i/item/9789240046979. Accessed: 2026.

[64] Bob Duncan and Mark Whittington. Creating an immutable database for secure cloud audit trail and system logging. In CLOUD COMPUTING 2017: The Eighth International Conference on Cloud Computing, GRIDs, and Virtualization, pages 54–59, Athens, Greece, 2017. IARIA. ISBN 978-1-61208-529-6.

[65] Daniel C Koboldt. Best practices for variant calling in clinical sequencing. Genome medicine, 12(1):91, 2020. doi: 10.1186/s13073-020-00791-w.

[66] Serena Dotolo, Riziero Esposito Abate, Cristin Roma, Davide Guido, Alessia Preziosi, Beatrice Tropea, Fernando Palluzzi, Luciano Giaco, and Nicola Normanno. Bioinformatics: from ngs data to biological complexity in variant detection and oncological clinical practice. Biomedicines, 10(9):2074, 2022. doi: 10.3390/biomedicines10092074.

[67] Alexei J Drummond, Marc A Suchard, Dong Xie, and Andrew Rambaut. Bayesian phylogenetics with beauti and the beast 1.7. Molecular biology and evolution, 29(8):1969–1973, 2012. doi: 10.1093/molbev/mss075.

[68] National Center for Biotechnology Information. NCBI submission portal. https://submit.ncbi.nlm.nih.gov/, 2024. Accessed: 2026.

[69] National Institute of Standards and Technology. Framework for improving critical infrastructure cybersecurity, version 1.1. Technical Report NIST.CSWP.04162018, U.S. Department of Commerce, National Institute of Standards and Technology, 2018.

[70] National Center for Biotechnology Information. table2asn: sequence submission preparation tool. https://www.ncbi.nlm.nih.gov/genbank/table2asn/, 2024. Accessed: 2026.

[71] Marc Ohm, Henrik Plate, Arnold Sykosch, and Michael Meier. Backstabber’s knife collection: A review of open source software supply chain attacks. In International Conference on Detection of Intrusions and Malware, and Vulnerability Assessment, pages 23–43. Springer, 2020. doi: 10.1007/978-3-030-52683-2_2.

[72] Vijay Singh Rathore, Nilanjan Dey, Vincenzo Piuri, Rosalina Babo, Zdzislaw Polkowski, and JMRS Tavares. Rising threats in expert applications and solutions. Advances in intelligent systems and computing, 1187, 2021.

